# Novel cholinesterase paralogs of *Schistosoma mansoni* have perceived roles in cholinergic signaling, glucose scavenging and drug detoxification and are essential for parasite survival

**DOI:** 10.1101/583658

**Authors:** Bemnet Tedla, Javier Sotillo, Darren Pickering, Ramon M. Eichenberger, Luke Becker, Alex Loukas, Mark S. Pearson

## Abstract

Cholinesterase (ChE) function in schistosomes is essential for orchestration of parasite neurotransmission but has been poorly defined with respect to the molecules responsible. Interrogation of the *S. mansoni* genome has revealed the presence of three ChE domain-containing genes (*Smche*)s, which we have shown to encode two functional acetylcholinesterases (AChE)s (*Smache1* – smp_154600 and *Smache3* – smp_136690) and a butyrylcholinesterase (BChE) (*Smbche1* – smp_125350). Antibodies to recombinant forms of each *Sm*ChE localized the proteins to the tegument and neuromusculature of adults and schistosomula and developmental expression profiling differed among the three molecules, suggestive of functions extending beyond traditional cholinergic signaling. For the first time in schistosomes, we identified ChE enzymatic activity in fluke excretory/secretory (ES) products and, using proteomic approaches, attributed this activity to the presence of *Sm*AChE1 and *Sm*BChE1. To address the hypothesis that tegumental AChE mediates exogenous glucose scavenging by the parasite, we show that RNAi-mediated knockdown of *smache1* and *smache3*, but not *smbche1*, significantly reduces glucose uptake by schistosomes. Parasite survival *in vitro* and *in vivo* was significantly impaired by silencing of each *smche*, either individually or in combination, attesting to the essential roles of these molecules. Lastly, in the first characterization study of a BChE from helminths, evidence is provided that *Sm*BChE1 may act as a bio-scavenger of AChE inhibitors as the addition of recombinant *Sm*BChE1 to parasite cultures mitigated the effect of the anti-schistosome AChE inhibitor DDVP (DDVP), whereas *smbche1*-silenced parasites displayed increased sensitivity to DDVP.

**Author summary:** Cholinesterases - aceytlcholinesterases (AChE)s and butyrylcholinesterases (BChE)s - are multi-functional enzymes that play a pivotal role in the nervous system of parasites by regulating neurotransmission through acetylcholine hydrolysis. Herein, we provide a detailed characterization of schistosome cholinesterases using molecular, enzymatic and gene-silencing approaches and show evidence for these molecules having roles in glucose scavenging and drug detoxification, in addition to their neuronal function. Further, we demonstrate the importance of these proteins to parasite development and survival through gene knockdown experiments in laboratory animals, providing evidence for the use of these proteins in the development of novel intervention strategies against schistosomiasis.

## Introduction

The functioning of the nervous system is a tightly regulated process controlled through multiple catalytic and non-catalytic signaling proteins. Among the catalytic molecules, cholinesterases (ChEs) play a pivotal role in regulating the signaling activity of the nervous system. There are two major types of ChEs, acetylcholinesterase (AChE) and pseudocholinesterase, or butyrylcholinesterase (BChE), and they can be distinguished both kinetically and pharmacologically [1]. AChE selectively hydrolyzes the neurotransmitter acetylcholine (ACh) to maintain neurotransmitter homeostasis [2] while the main role of BChE is widely accepted to be the detoxification of organophosphorus esters which are inhibitors of AChE [3]. ChEs are generally believed to be functionally redundant in cholinergic signaling with the main differences between paralogs lying in their spatial and temporal expression as well as non-cholinergic functionality [4, 5].

The nervous system of helminths has long been a potential target for therapeutic agents as it plays several crucial roles in parasite biology that are fundamental to survival, including coordinating motility within and outside of the host, feeding and reproduction [6–10]. The *Schistosoma* nervous system is particularly important in this respect as this parasite lacks a body cavity and circulating body fluid [11, 12] and, as a result, its signaling functions are chiefly achieved via neurotransmission. The primary neurotransmitter that schistosomes utilize is acetylcholine (ACh), which allows muscle contraction. The physiological concentration of ACh, however, must be maintained otherwise it triggers paralysis and this is achieved primarily through the action of AChE [6–8].

While AChE activity has been documented extensively in *S. mansoni* (reviewed in [13]), most of the work has involved studies on parasite extracts or native *Sm*ChE purified by inhibitor-affinity chromatography, making it difficult to attribute function to any one particular *Sm*ChE molecule. Further, more recent, but still “pre-genomic”, studies have documented only one AChE-encoding gene in *S. mansoni* and other species [14, 15]. In 2016, You *et al*. characterized AChE activity in *S. japonicum* extracts and at a molecular level, but only through the expression of one recombinant AChE [16]. Moreover, to the best of our knowledge, genes encoding proteins with BChE activity have not been previously described in schistosomes or any other helminth. Interrogation of the now fully annotated *S. mansoni* genome [17] has revealed three different *Sm*ChE paralogs; however, their individual contributions to ChE function remain unknown.

Traditionally, ChEs have been regarded solely as neurotransmitter terminators; however, there is increasing data to suggest that these enzymes play a variety of roles that extend beyond this cholinergic function due to their presence in multiple cell types and subcellular locations [4, 5, 18, 19]. In schistosomes, AChE has been localized to the tegument as well as the neuromusculature [16, 20, 21] and a proposed function for tegumental AChE has been mediation of glucose uptake by the parasite from the external environment [22]. The exact mechanism for this process is still unclear but the proposed initiating step is by limiting the interaction of host ACh with tegumental nicotinic ACh receptors (nAChRs), a hypothesis bolstered by the observation that glucose uptake is ablated by the use of membrane-impermeable AChE and nAChR inhibitors in *S. mansoni* [22, 23] and RNAi-mediated AChE silencing in *S. japonicum* [24]. The nAChRs are also associated both spatially and temporally with surface AChE expression and are concentrated on the tegument [25], the major site of glucose uptake [26].

Many intestinal nematodes secrete AChE [27–30], which, where studied, orchestrate exogenous cholinergic activities. It has also been indirectly shown that the nematode *N. brasiliensis* employs parasite-derived AChE to alter the host cytokine environment to inhibit M2 macrophage recruitment, a condition favorable to worm survival [31]. Despite this breadth of literature in nematodes, there has been no documentation of secreted AChE activity from schistosomes.

Herein we describe and functionally characterize using gene silencing and enzymatic approaches, a novel AChE and BChE from *S. mansoni* and further characterize the only previously identified AChE-encoding gene from the parasite. Importantly, we show through gene knockdown that each *smche* is essential to *S. mansoni* development and survival, highlighting them as targets for novel anti-schistosomal intervention strategies.

## Results

### Identification of novel genes encoding ChE proteins in S. mansoni

Three putative ChE paralogs were identified from interrogation of the *S. mansoni* genome: *smache1* (Smp_154600), *smbche1* (Smp_125350) and *smache3* (Smp_136690). The predicted *Sm*ChEs were then aligned with characterized AChE enzymes from *Homo sapiens*, the electric eel *Torpedo californica*, and the nematode *Caenorhabditis elegans* (Figure 1). Homology analysis of amino acid sequences revealed that *Sm*AChE1, *Sm*BChE1, and *Sm*AChE3 share (32-35%) sequence identity and (49-52%) sequence similarity. Further, all *Sm*ChEs have 36-40% amino acid identity with *H. sapiens* and *T. californica* AChE. All identified *Sm*ChEs had ChE-specific characteristics, including a catalytic triad with an active site serine, which is required for ester hydrolysis [32]. Interestingly, the His residue of the catalytic triad of *Sm*BChE1 appears to have been substituted for Gln, a change consistent among all the BChE1 homologs shown for other Platyhelminthes, but not nematode or model organism BChE1 sequences (Figure S1). A 3D model of the three *Sm*ChEs was constructed by homology modeling with AChE from model organisms (*H. sapiens* and *T. californica* (Figure S2). All three *Sm*ChEs exhibited predicted folding characteristics of the functional globular enzymes as most of the α-helical and β-stranded sheets were tightly aligned. Each predicted *Sm*ChE structure consisted of a ChE catalytic domain but, although the core architecture of the catalytic gorges was well aligned, regions that are associated with substrate specificity and catalytic efficiency were disparate. In particular, and in agreement with the sequence alignment, the catalytic triad of *Sm*BChE1 was predicted to be Ser-Gln-Glu instead of the canonical Ser-His-Glu present in the other two paralogs. A phylogenetic tree of the alignment (Figure S3) shows that *Sm*ChEs were clustered into three distinct branches, with *Sm*ChE1 being phylogenetically distinct from *Sm*BChE1 and *Sm*AChE3. In addition, each *Sm*ChE was grouped together with closely related flatworms, including other *Schistosoma* species.

**Figure 1.**
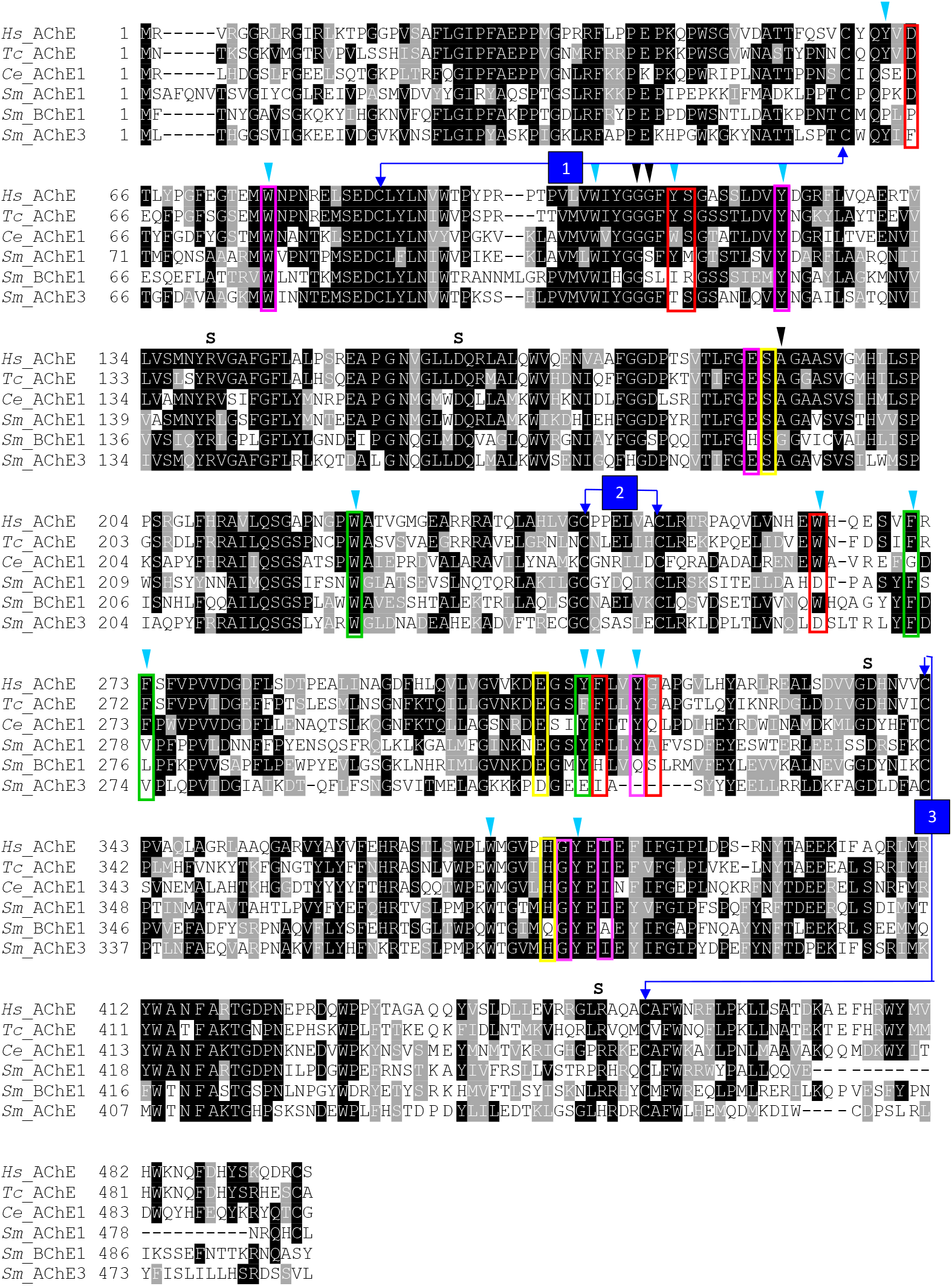
The amino acid sequence alignment and phylogeny of ChEs from *S. mansoni* and other species. Light blue arrowheads = the 14 aromatic rings, black arrowheads = oxyanion holes, S= salt bridges, red boxes= PAS, yellow boxes = catalytic triad, green boxes = acyl binding pocket, numbered arrows = disulfide bonds and magenta box = peripheral anionic site. Accession numbers: *H. sapiens* (NP000656), *T. californica* (CAA27169), *C. elegans* (NP510660), *Sm*AChE1 (Smp_154600), *Sm*BChE1 (Smp_125350), *Sm*AChE3 (Smp_136690).

Importantly, as shown in the sequence alignment, *Sm*ChEs are divergent from the human homolog. Reflective of the catalytic triad residue difference (Figure S1), trematode BChEs are phylogenetically divergent from nematode and human BChEs (Figure S4).

### Developmental expression analysis of SmChE genes

Gene expression patterns of the three *Sm*ChE paralogs across different developmental stages were measured using semi-quantitative qPCR (Figures 2A-C) and this data was used to generate a comparative expression heat map of all three genes (Figure 2D). While all *smche* developmental expression patterns were variable, the transcript levels of all three genes were relatively lower in cercariae compared to the other developmental stages. Overall, the transcript levels of *smache1* and *smache3* genes in most life stages were higher than that of *smbche1*. In adult worms, *smache1* was expressed at higher levels, specifically in male parasites, followed by sporocysts.

**Figure 2.**
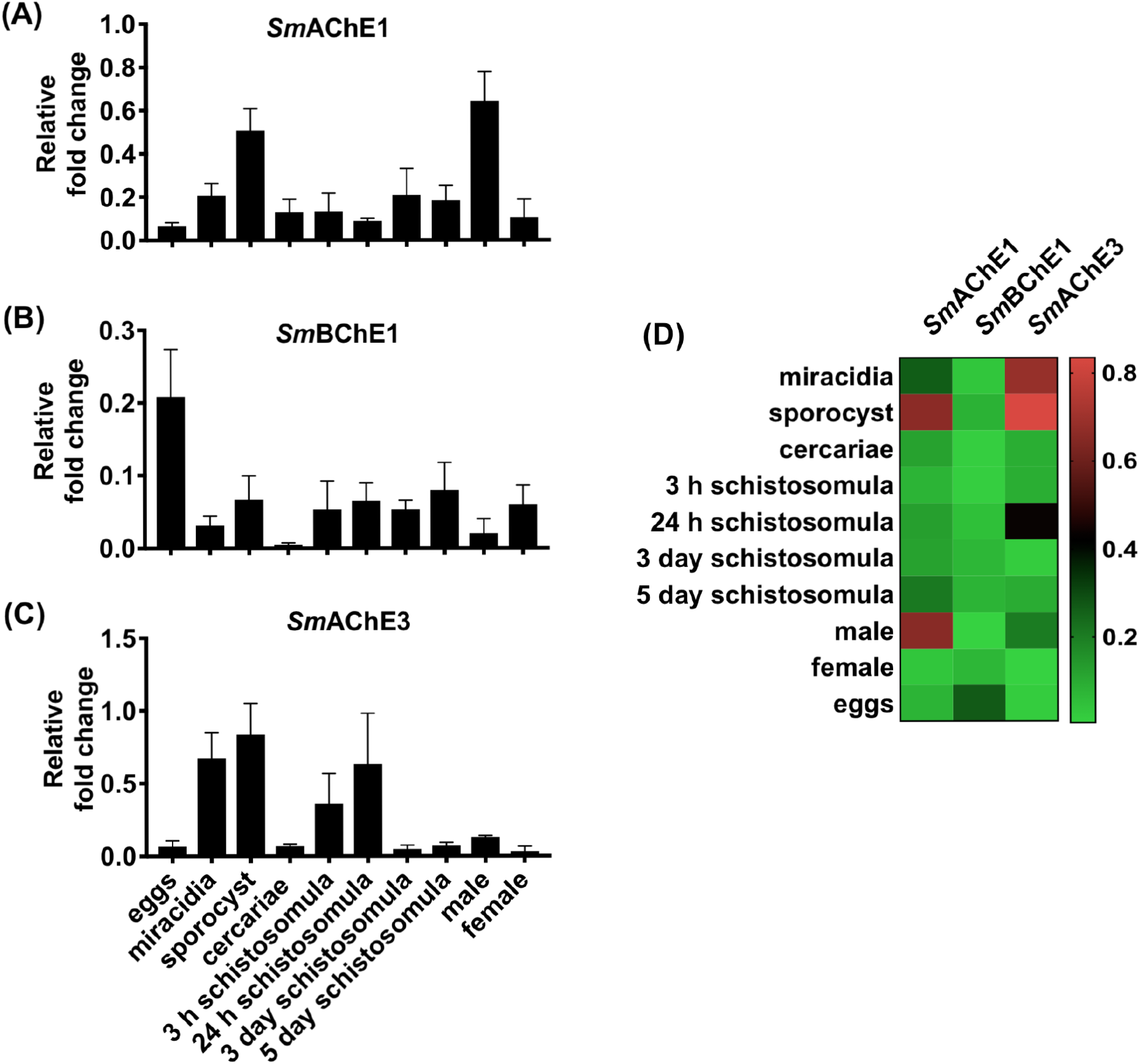
Developmental expression profiles of *smache1, smbche1* and *smache3*. The expression of (A) *smache1*, (B) *smbche1* and (C) *smache3* genes at different developmental stages of *S. mansoni* as quantified by qPCR analysis. (D) The heat map shows the comparative expression pattern of the paralogs in each developmental stage. Data are presented as mean ± SEM of five independent experiments and are normalized to the *smcox1* housekeeping gene.

### Immunolocalization of SmChEs

To gain insight into the anatomical sites of expression of ChE proteins in *S. mansoni, Sm*ChEs were immunolocalized in whole juvenile and sectioned adult parasites. In adults, and consistent with their predicted cholinergic function, all *Sm*ChEs were expressed throughout the worms’ internal structures (presumably localizing to the neuromusculature) and on the tegument surface. *Sm*AChE1 was the least uniformly distributed of all *Sm*ChEs, localizing mostly to the tegument (Figure 3A). Additionally, anti-*Sm*ChE antibodies were able to detect homologous ChEs in adult *S. haematobium* sections. *Sm*ChE proteins were detected in all stages of larval development tested and, as was the case with adult worms, localized to the tegument (Figure 3B).

**Figure 3.**
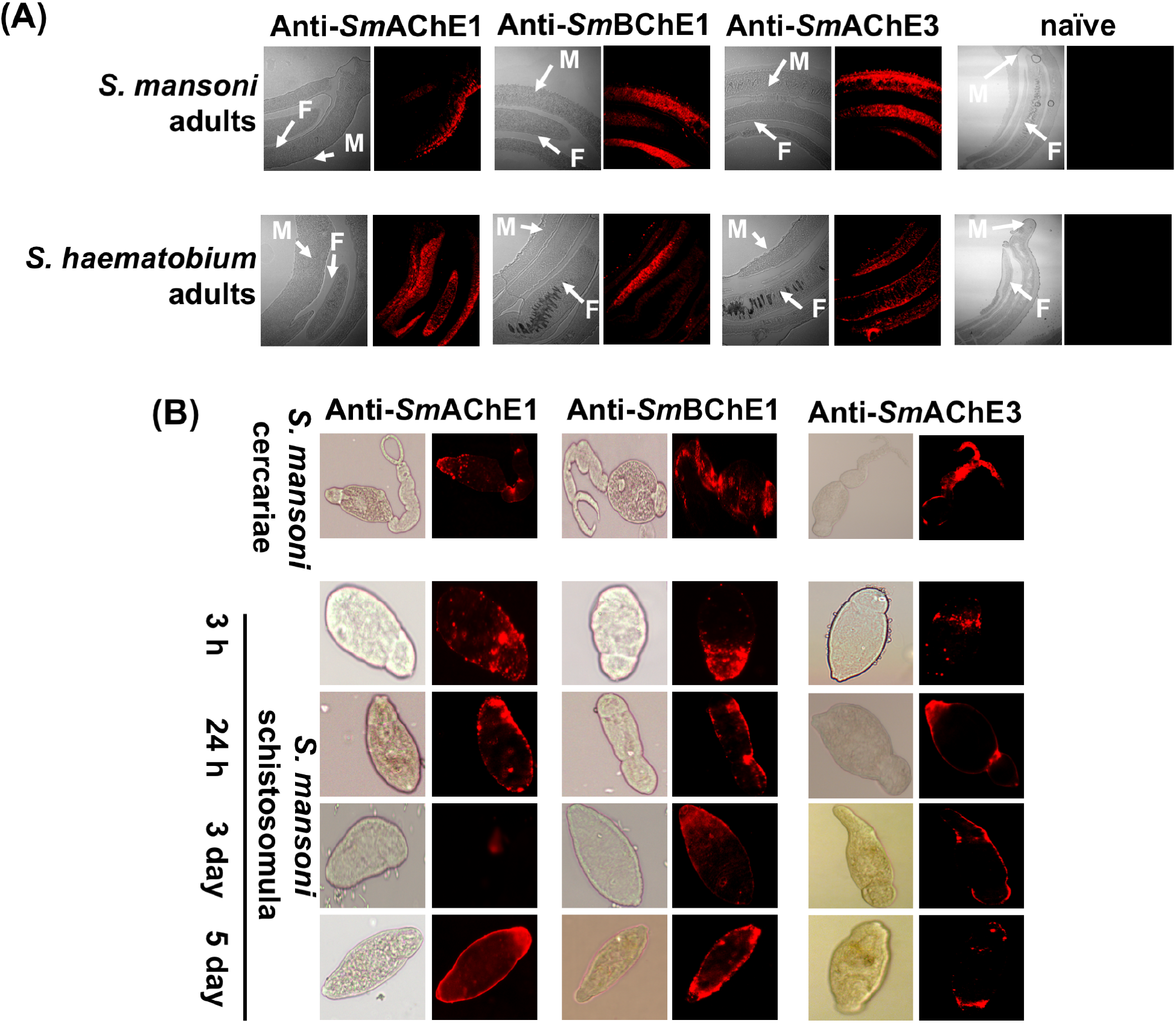
Immunofluorescent localization of *Sm*ChEs. Fluorescence and brightfield images of (A) male (M) and female (F) *S. mansoni* and *S. haematobium* adult worm sections. (B) Live, fixed cercariae, and schistosomula at 3 h, 24 h, 3 days and 5 days after transformation. Both adult sections and juvenile parasites were labeled with either anti-*Sm*AChE1, anti-*Sm*BChE1 or anti-*Sm*AChE3 primary antibody (1:100 in PBST) followed by goat-anti-mouse IgG-alexafluor 647 (1:200 in PBST). Naive mouse sera was used as a negative control.

### Expression and ChE activity of fSmChEs

Soluble, functionally active proteins were expressed in *P. pastoris*, purified via IMAC and tested for ChE activity. Both f*Sm*AChE1 and f*Sm*AChE3 demonstrated significantly stronger hydrolase activity when AcSCh was used as a substrate, compared to f*Sm*BChE1 and, conversely, f*Sm*BChE1 hydrolyzed BcSCh to significantly higher levels compared to f*Sm*AChE1 and f*Sm*AChE3 (Figure 4A). All paralogs exhibited Michaelis–Menten kinetics (Table 1) when hydrolyzing their designated substrate, with f*Sm*AChE1 having a substrate affinity approximately twice that of f*Sm*AChE3. In addition, preferred substrate activity of both f*Sm*AChE1 and f*Sm*AChE3 was inhibited by DDVP, an AChE inhibitor, whereas iso-OMPA, a specific inhibitor of BChE, only inhibited *Sm*BChE1 activity (Figure 4B).

**Figure 4.**
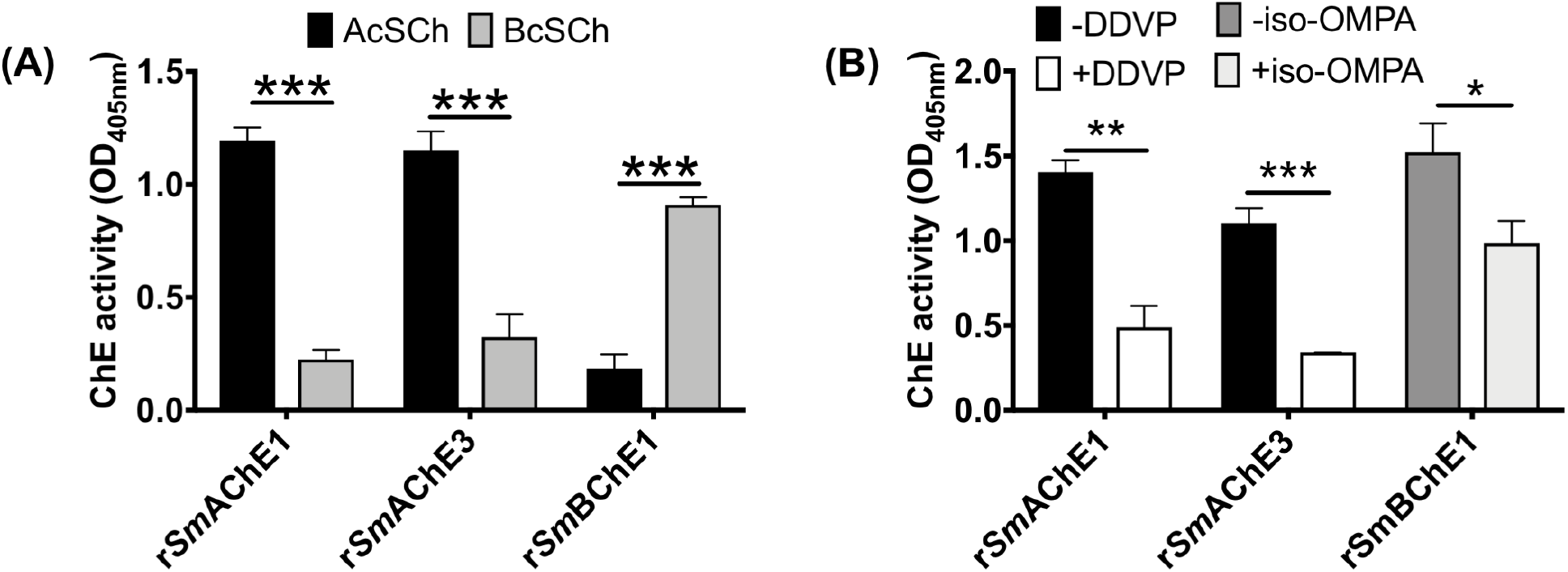
Enzymatic activity of f*Sm*ChEs. (A) Cholinergic substrate preference (AcSCh or BcSCh) of each f*Sm*ChE. (B) Inhibition of f*Sm*AChE1 and f*Sm*AChE3 with DDVP (AcSCh used as a substrate) and inhibition of f*Sm*BChE1 with iso-OMPA (BcSCh used as a substrate). Data are presented as mean ± SEM of triplicate experiments and differences between groups were measured by the student’s *t* test. \**P* ≤ 0.05, \*\**P* ≤ 0.01, \*\*\**P* ≤ 0.001.

**Table 1.** Kinetic parameters of f*Sm*ChEs

| SmChE paralog | V<sub>max</sub> (nmol/min/mg) | K<sub>m</sub> (mM) |
| --- | --- | --- |
| fSmAChE1 | 5.57 ± 0.54* | 5.83 ± 1.62* |
| fSmAChE3 | 5.59 ± 1.37* | 10.87 ± 6.17* |
| fSmBChE1 | 1.7 ± 0.09<sup>#</sup> | 34.38 ± 12.71<sup>#</sup> |

\*Hydrolysis of AcSCh.
<sup>#</sup>Hydrolysis of BcSCh.

### BChE and secreted ChE activity in schistosomes

Although the presence of nonspecific ChE activity has long been known in schistosomes [33], the identity of the gene product and its function remain unknown. Prompted by the identification of *Sm*BChE1 as a BChE, based on its substrate preference and enzymatic inhibition by iso-OMPA, we sought to investigate the distribution of BChE activity in juvenile and adult schistosomes. Extracts from *S. mansoni* schistosomula had higher BChE activity compared to *S. mansoni* adult worms (Figure 5A), and that activity was significantly greater in *S. mansoni* compared with *S. haematobium* adults (Figure 5B). Varied amounts of AChE activity were detected in ES from all developmental stages tested. ES products from adult males had double the AChE activity of adult female ES products (*P* < 0.001), while cercariae ES exhibited the highest activity (at least ten-fold more than male ES products (*P* < 0.0001)) and egg ES had the lowest (Figure 5C). Availability of ES precluded the measurement of secreted BChE activity from all developmental stages but, of those tested, activity in schistosomula ES products was the highest - twice as high as that of adult (*P* < 0.01) and cercariae (*P* < 0.01) ES (Figure 5D). *Sm*ChEs were purified from ES products of *S. mansoni* adult worms using edrophonium–sepharose affinity chromatography. Purification resulted in an activity increase of more than 200-fold relative to crude ES (Figure 5E). Resolution of the purified proteins by SDS-PAGE resulted in a doublet with a major band migrating at 70 kDa under denaturing and reducing conditions (Figure 5F). The identity of purified, secreted *Sm*ChEs was substantiated by in-gel LC-MS/MS analysis with the peptide data generated used to interrogate the *S. mansoni* proteome (predicted from the *S. mansoni* genome - http://www.genedb.org/Homepage/Smansoni). The false discovery rate was set at <1% and only proteins with at least two unique peptides having significant Mascot identification scores (*P* < 0.05) were considered. The top protein hits were identified as *Sm*AChE1 (Smp_154600) and *Sm*BChE1 (Smp_125350); *Sm*AChE1 had a relative abundance of more than 40-fold that of *Sm*BChE1 (supplementary table 3).

**Figure 5.**
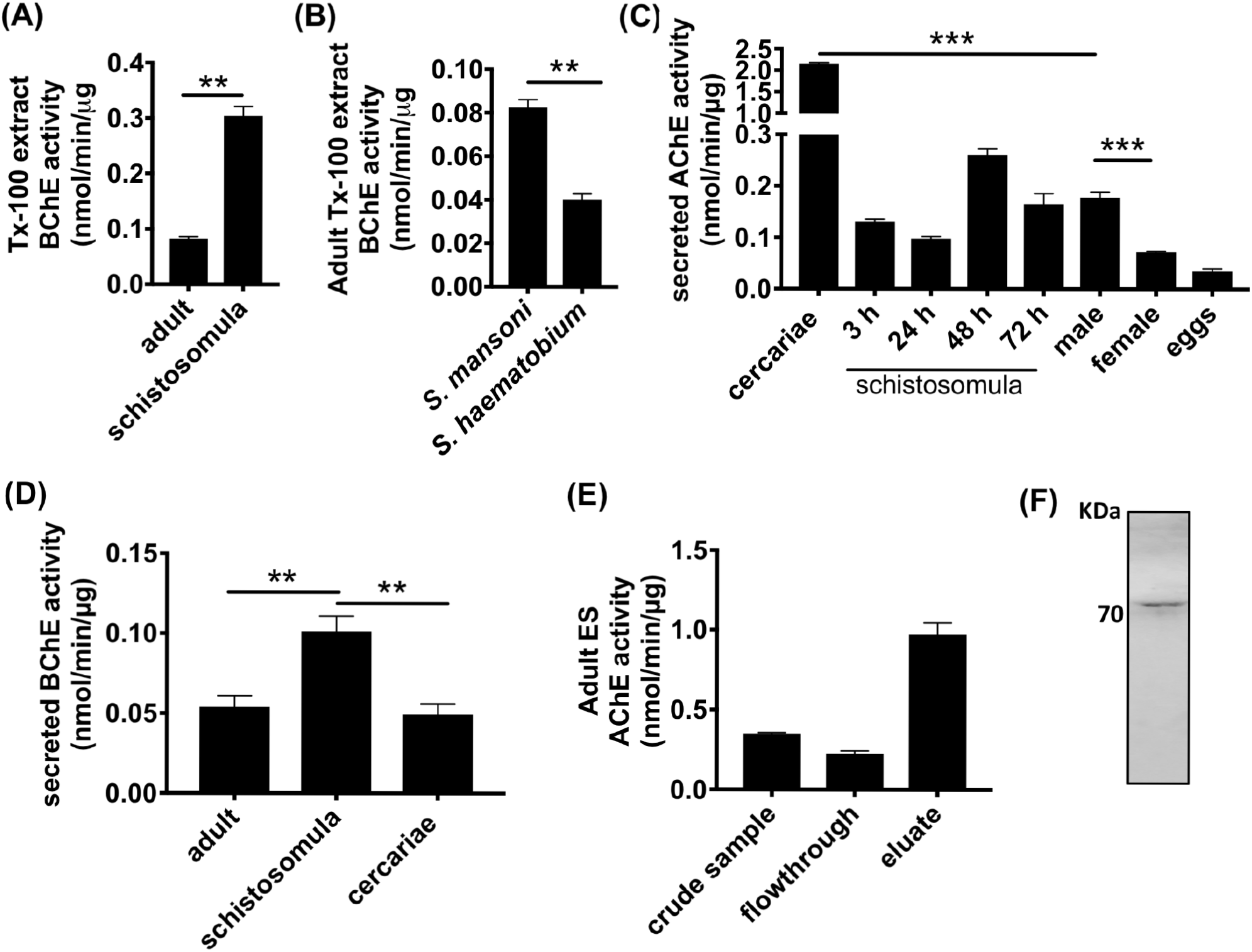
BChE and secreted ChE activity in schistosomes. (A) BChE activity in *S. mansoni* adults and schistosomula TX-100 extracts. (B) BChE activity in TX-100 extracts from *S. mansoni* and *S. haematobium*. (C) AChE and (D) BChE activity of ES products from different developmental stages of *S. mansoni*. (E) AChE activity and (F) SDS-PAGE analysis of purified, secreted *Sm*ChEs. Data are presented as mean ± SEM of triplicate experiments and differences between groups were measured by the student’s *t* test. \**P* ≤ 0.05, \*\**P* ≤ 0.01.

### RNAi-mediated smche transcript and SmChE protein reduction

Schistosomula electroporated with *smache1* siRNA showed respective decreases in *Sm*AChE1 mRNA levels of 55.4% (*P* ≤ 0.05) and 81.3% (*P* ≤ 0.001) at one and seven days post-treatment, respectively, compared to the *luc* control (Figure 6A), while treating parasites with *smbche1* siRNA caused 32.0% (*P* ≤ 0.001) and 84.5% (*P* ≤ 0.001) suppression of *smbche1* mRNA expression at day 3 and 7 after electroporation, respectively, compared to the *luc* control (Figure 6B). Treatment of schistosomula with *smache3* siRNA resulted in respective decreases in *smache3* mRNA levels of 27.4% (*P* ≤ 0.001) and 47.2% (*P* ≤ 0.01) three and seven days after electroporation, compared to the *luc* control (Figure 6C). Schistosomula electroporated with a cocktail of all three *smche* siRNAs showed decreases of all three transcript levels over time, with *smache3* mRNA levels decreasing by an average of 90% (*P* ≤ 0.001) by day 3 after treatment, compared to the *luc* control (Figure 6D).

**Figure 6.**
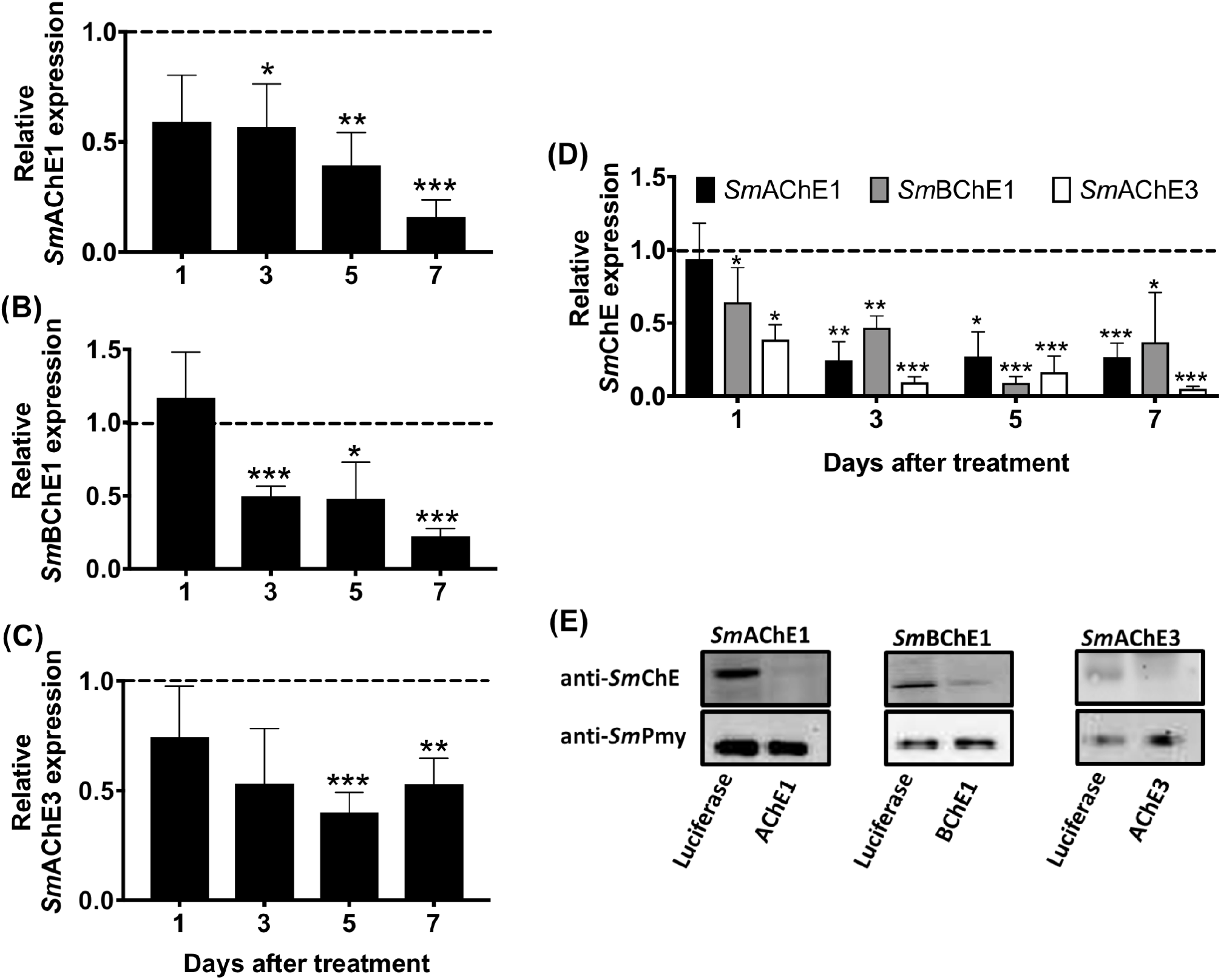
Suppression of *smche* mRNA transcript and protein expression in schistosomula by RNAi. Individual siRNA treatment with (A) *smache1*, (B) *smbche1* or (C) *smache3* siRNAs. (D) Treatment with a cocktail of *smache1, smbche1* and *smache3* siRNAs. Transcript levels of each *smche* in parasites treated with *smche* siRNAs are shown relative to *smche* transcript expression in schistosomula treated with the *luc* control siRNA (dashed line) and represent the mean ± SEM of triplicate qPCR assays from 2 biological replicates of each treatment. Transcript expression in all parasites was normalized with the housekeeping gene, *smcox1*. Differences in transcript levels (relative to the *luc* control) were measured by the student’s *t* test. \**P* ≤ 0.05, \*\**P* ≤ 0.01, \*\*\**P* ≤ 0.001. (E) Western blot of day 7 schistosomula extracts following treatment with *smche* or *luc* siRNAs. Extracts were immunoblotted with the corresponding anti-*Sm*AChE1, anti-*Sm*BChE or anti-*Sm*AChE3 polyclonal antibody. An antibody against *Sm*Pmy (paramyosin) was used as a loading control.

Seven days after treatment with *Sm*AChE1, *Sm*BChE1 or *Sm*AChE3 siRNAs, schistosomula showed decreases in *Sm*AChE1, *Sm*BChE1 or *Sm*AChE3 protein expression of 73%, 59% and 46%, respectively, compared to luciferase controls (Figure 6E).

### Suppression of SmChE activity

Suppression of AChE activity was seen in *smache1* and *smache3* siRNA-treated parasites from 5 and 3 days after electroporation, respectively (Figure 7A), compared to the *luc* control, while schistosomula treated with *smbche1* siRNA did not show any significant reduction in BChE activity, even 7 days after electroporation (Figure 7B). Parasites electroporated with a cocktail of all three *smche* siRNAs showed significant decreases in AChE activity at 3 days (62% reduction, *P* ≤ 0.001), 5 days (67% reduction, *P* ≤ 0.001) and 7 days (71% reduction, *P* ≤ 0.001) after treatment (Figure 7C). BChE activity was not measured in the cocktail siRNA treatment group.

**Figure 7.**
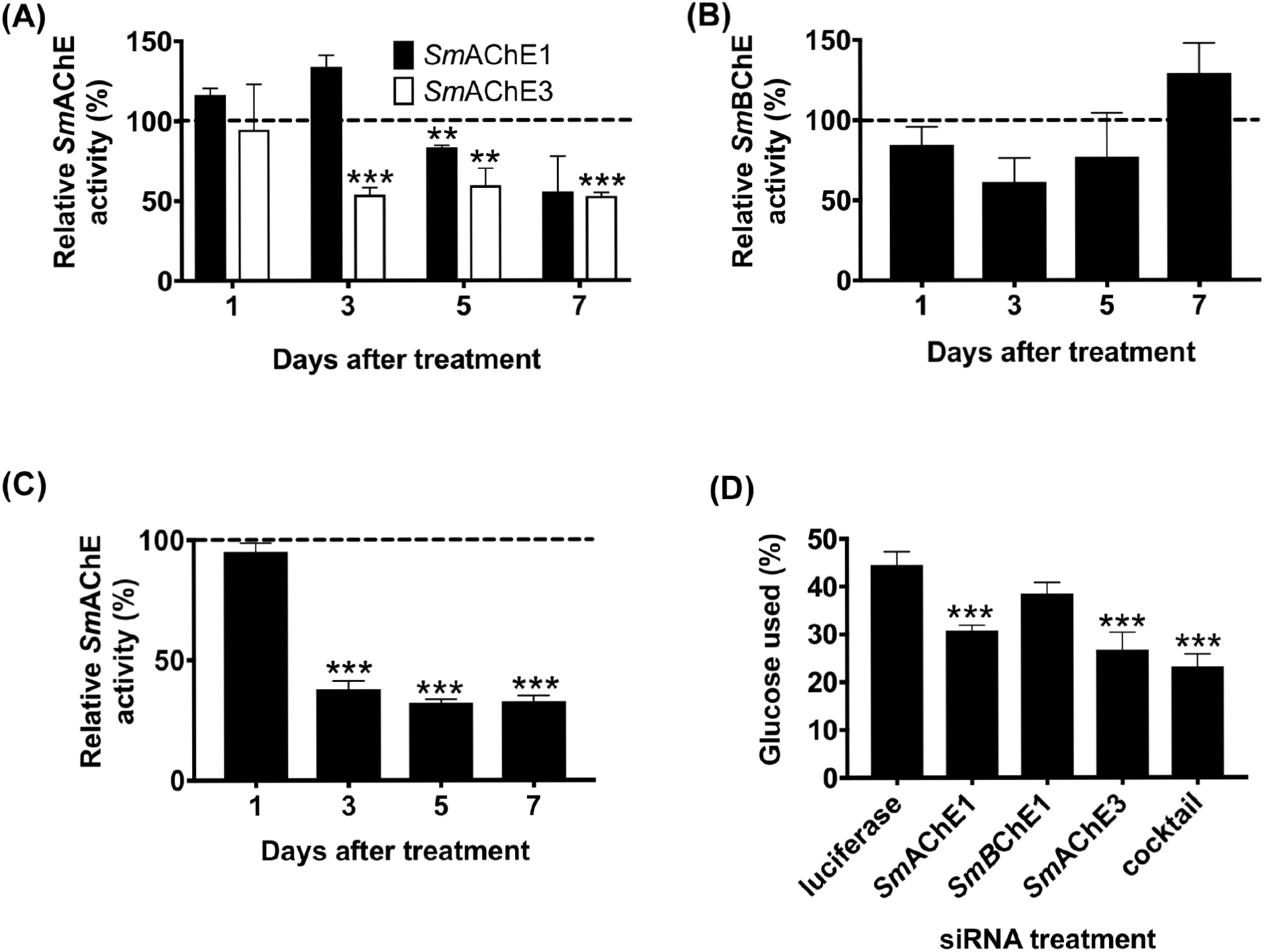
Effects of *smche* knockdown on cholinesterase activity and glucose uptake. (A) AChE activity of extracts from schistosomula treated with *smache1, smache3* or *luc* siRNAs (dashed line). (B) BChE activity of extracts from schistosomula treated with *smbche1* or *luc* siRNAs (dashed line). (C) AChE activity of extracts from schistosomula treated with all 3 siRNAs or *luc* siRNA (dashed line). (D) Glucose uptake by schistosomula 48 h after treatment with *smche* siRNAs. Schistosomula (5 days old – 5,000/treatment) were electroporated with either *luc* or *smche* siRNAs and glucose consumption was measured 48 h after treatment. Data represents mean ± SEM of duplicate assays from 2 biological replicates of each treatment. Differences (relative to the *luc* control) were measured by the student’s *t* test. \*\**P* ≤ 0.01, \*\*\**P* ≤ 0.001.

Individual silencing of *smache1* or *smache3* genes and combined silencing of all three *smche* genes reduced glucose uptake in schistosomula by 24.9% (*P* ≤ 0.001), 32.34% (*P* ≤ 0.001) and 38.61% (*P* ≤ 0.001) at 48 h post-treatment, respectively, relative to the *luc* control. However, *smbche1*-silenced parasites showed no significant changes in glucose uptake at the same timepoint and there was no difference in the glucose consumed by the *smache1* or *smache3* siRNA-treated groups compared with the cocktail siRNA-treated group (Figure 7D). Transcript levels of the glucose transporters, *sgtp1* and *sgtp4*, were neither decreased nor significantly increased in individual or cocktail *smche*-silenced parasites (Figure S5).

### Effects of smche silencing on schistosomula viability in vitro and development in vivo

Parasites treated with *smache1, smbche1* or *smache3* siRNAs showed significant decreases in viability at days 3, 5 and 1 after treatment, respectively, compared to *luc* controls. At days 5 and 7 post-treatment, the most significant decrease in parasite viability was seen in the group which received the cocktail siRNA treatment, compared to *luc* controls. Furthermore, viability in this group was also significantly lower than it was for any individual treatment at these two time points (Figure 8A).

**Figure 8.**
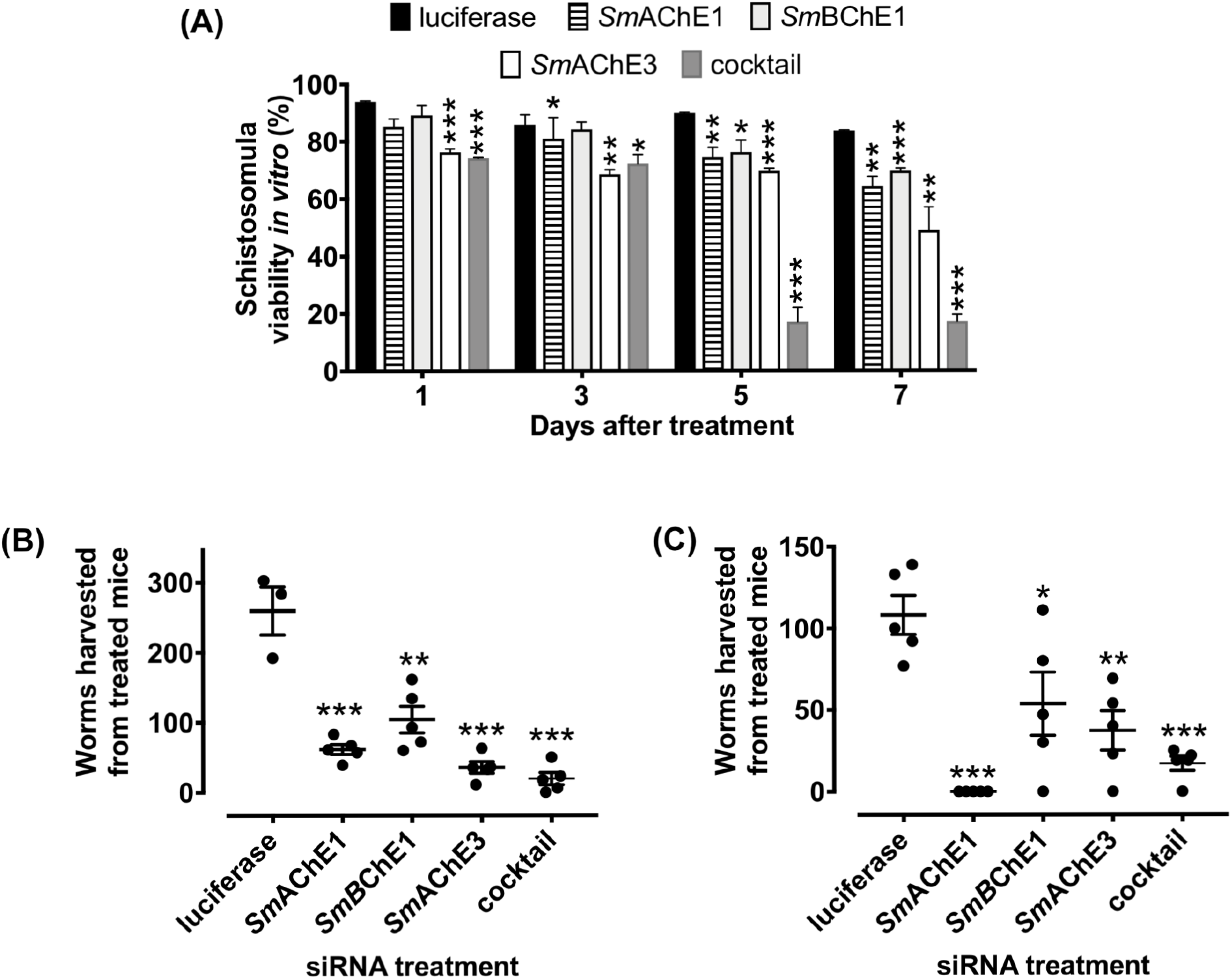
Effects of *smche* silencing on schistosomula viability *in vitro* and development *in vivo*. (A) Schistosomula treated with individual or a cocktail of all 3 *smche* siRNAs or *luc* siRNA were cultured for 7 days in complete Basch medium with viability determined at day 1, 3, 5 and 7 after treatment by Trypan Blue exclusion (mean ± SEM of duplicate assays from 2 biological replicates of each treatment). (B and C) One-day-old schistosomula treated with individual or a cocktail of all three *smche* siRNAs or *luc* siRNA were intramuscularly injected (2,000 parasites) into mice. After 3 weeks, adult worms were recovered and counted. Data from two independent experiments are shown. Differences between *smche*- and *luc*-treated groups were measured by the student’s *t* test. \**P* ≤ 0.05, \*\**P* ≤ 0.01, \*\*\**P* ≤ 0.001.

To examine whether RNAi-mediated *smche* suppression reduced parasite viability *in vivo*, mice were infected with *smche*-silenced parasites and worm burdens were measured after three weeks. From two independent experiments, there was an average 88.15%, 55.15%, 75.95% and 88.35% decrease in adult fluke burdens from mice injected with *smache1-, smbche1-, smache3*- and *smche* cocktail-silenced schistosomula, respectively, compared to mice infected with *luc*-treated parasites (Figures 8B and C). All worm burden decreases were significant and there was no significant difference in fluke burdens between mice injected with *luc*-treated parasites and non-electroporated control parasites. All mice had been successfully infected with parasites, as serum from necropsied mice contained parasite-specific antibodies (data not shown). Compared to *luc*-treated parasites, worms recovered from *smche*-treated parasites showed no difference in *smche* transcript levels (data not shown).

### Bio-scavenging of carboxylic esters by SmBChE1

The hypothesis that *Sm*BChE may act as a molecular decoy in schistosomes and detoxify the effects of organophosphorus AChE inhibitors was examined by testing whether (a) inhibition of parasite-derived BChE potentiated the effects of DDVP (an organophosphorous AChE inhibitor) and (b) addition of exogenous BChE (f*Sm*BChE1) mitigated the effects of DDVP. DDVP activity in schistosome extracts significantly increased in the presence of increasing amounts of the BChE inhibitor, iso-OMPA (Figure 9A) and DDVP-mediated killing of schistosomula was significantly increased in the presence of iso-OMPA (60.9% compared with 21.8%; *P* < 0.0001) (Figure 9B). Further, *smbche1*-silenced schistosomula were significantly more susceptible to DDVP-mediated killing than *luc*-treated controls (83.44% compared with 22.95%; *P* < 0.0001) (Figure 9C). Conversely, DDVP-induced inhibition of AChE in extracts was completely ablated in the presence of f*Sm*BChE (Figure 9D) and schistosomula were increasingly resistant to DDVP-mediated killing with the addition of increasing amounts of recombinant protein to the culture media (Figure 9E).

**Figure 9.**
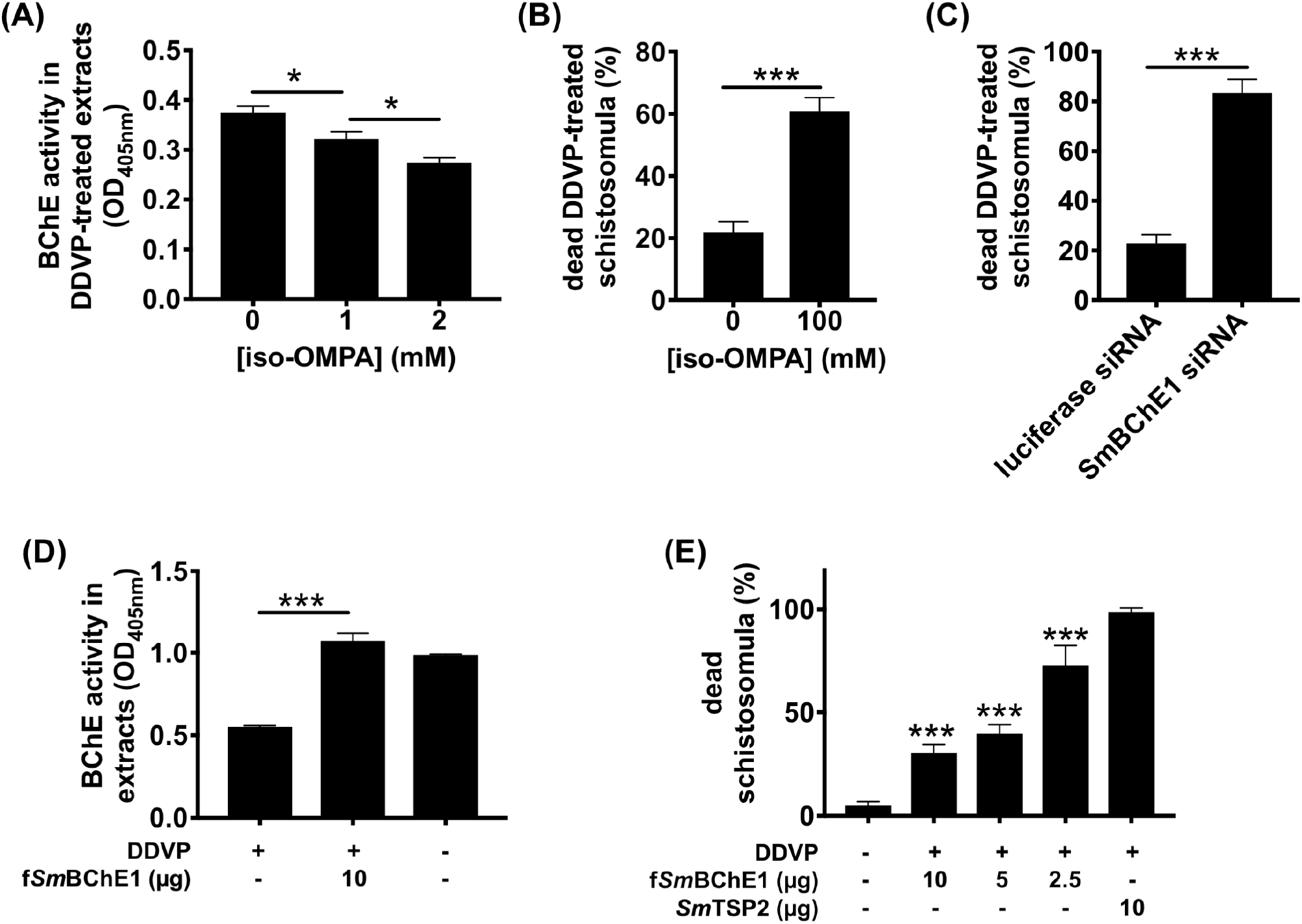
*Sm*BChE1 bio-scavenges DDVP and protects parasites against DDVP-induced effects. (A) Schistosomula extracts were treated with DDVP (1 μM), or pretreated with iso-OMPA (1 and 2 mM) and then DDVP, before assaying AChE activity. (B) Schistosomula were treated with DDVP (1 μM) or pretreated with iso-OMPA (100 μM) and then DDVP, and parasite viability was measured 5 h after treatment. (C) *smbche1*-silenced or *luc* siRNA-treated schistosomula were treated with DDVP (1 μM) and parasite viability was measured 5 h after treatment. (D) Schistosomula extracts were pre-incubated with f*Sm*BChE1 (10 μg) then treated with DDVP (1 μM), or treated with DDVP alone, before assaying AChE activity. (E) DDVP was pre-incubated with f*Sm*BChE1 (10, 5 and 2.5 μg) or 10 μg of *Sm*TSP2 for 1 h before being used to treat schistosomula. Parasite viability was measured 24 h post-treatment. For all assays, data are the average of triplicate biological and technical experiments ± SEM and differences were measured by the student’s *t* test. \**P* ≤ 0.05, \*\*\**P* ≤ 0.001, \*\*\*\**P* ≤ 0.0001.

## Discussion

Cholinesterase (ChE) activity in *S. mansoni* was first described by Bueding in 1952 [33] and was well characterized biochemically in the four decades succeeding this discovery. The technological limitations of this time period meant that most of the evidence for *Sm*ChEs came from whole worm studies and analyses of crude parasite extracts (reviewed in [13]), which could not ascribe ChE activity to any particular protein. Several studies in the early 2000’s characterized a single AChE from *S. mansoni* (Smp_154600 in the current gene annotation nomenclature) and its direct homolog in other species of schistosomes [14, 15, 21], but lack of a comprehensive schistosome genome annotation at the time precluded identification of more ChE family members. Interrogation of the most recent iteration of the *S. mansoni* genome assembly has identified two additional ChE-encoding genes that are paralogs to Smp_154600 (which we have termed *Sm*AChE1); Smp_125350 (*Sm*BChE1) and Smp_136690 (*Sm*AChE3). In this current study, we have provided a more in-depth characterization of the previously documented *Sm*AChE1 and described two novel ChEs from *S. mansoni: Sm*AChE3 – an AChE not previously reported; and *Sm*BChE1 – a BChE which, to the best of our knowledge, has never been documented in the helminth literature.

All *Sm*ChEs share a modest level of identity which is consistent with their divergence over evolutionary time, an occurrence that is possibly due to a series of gene duplications due to the phylogenetic distance between the relative clades. This divergence between *Sm*ChEs and, also, ChEs of other organisms, provides evidence for the increasing reports of non-cholinergic functions of ChEs in the literature. Additionally, the relative lack of sequence identity between *Sm*ChEs and human ChEs suggests potential scope for the development of intervention strategies targeting schistosome ChEs that will not affect the host. Despite the diversity between ChEs, all enzymes analyzed herein would appear to be enzymatically active as they possessed a catalytic triad with an active site serine, the amino acid responsible for ester hydrolysis [32]. It is interesting to note, however, the catalytic triad His – Gln substitution in *Sm*BChE1 (and the other platyhelminth BChE1 homologs); while this change is not a hallmark of model BChEs, that it occurs within an entire parasite lineage is noteworthy and will be investigated further.

The transcript levels of each *smche* varied among parasite developmental stages and this is likely a response to the differing cholinergic and cholinesterase-independent requirements of the parasite throughout its lifecycle. For example *smache1* is expressed at a higher level in adult males than females, probably due to the more “muscular” roles of attachment and movement orchestrated by the male compared to the female, which remains sedentary once inside the gynecophoric canal of the male [34]. Expression of *smbche1* was highest in the egg stage; there is evidence for BChE involvement in chicken embryo neurogenesis and development, independent of its enzymatic function [35], which suggests that *Sm*BChE1 could play a role in parasite embryogenesis. The miracidium and sporocyst stages had the highest levels of *smache3* expression, in agreement with Parker-Manuel et al [36].

Immunolocalization of the *Sm*ChEs revealed expression in the neuromusculature and tegument to varying degrees, depending on the paralog, and is consistent with early localization experiments [20], although the antibodies used in those studies were raised against AChEs purified from parasite extracts and so the localization could not be attributed to a specific family member. Localization to the neuromusculature relates to the proteins’ traditional cholinergic functions whereas tegumental distribution is suggestive of non-neuronal cholinergic and/or non-cholinergic roles. Indeed, surface-expressed *Sm*AChE has been implicated in mediating glucose scavenging by the parasite, as this process can be ablated by membrane-impermeable AChE inhibitors [22, 23]. Tegumental *Sm*AChE may also act to hydrolyze exogenous ACh, neutralizing its immune-mediating function to create an environment more conducive to parasite establishment [31]. The localization of these proteins to the tegument of schistosomula should also be noted since early developing schistosomula are considered most vulnerable to immune attack [37], and so *Sm*AChE-targeted immunotherapeutics could be used effectively to vaccinate against schistosomiasis. Indeed, antibodies against *Sm*AChEs have been shown to interact with the surface of schistosomula, resulting in complement-dependent killing of the parasite [38].

Full-length and functional *Sm*ChEs were expressed in *P. pastoris. Sm*AChE1 had preferred substrate specificity for AcSCh over BcSCh, albeit at a three-fold lower affinity than previously reported for *Sm*AChE1 expressed in *Xenopus laevis* oocytes [15]. *Sm*AChE3 also had a substrate preference for AcSCh and an affinity twice that of *Sm*AChE1. Extremely low enzyme activity was observed with *Sm*BChE1 when AcSCh was used as a substrate, but enzymatic activity significantly increased with the use of BcSCh as the substrate. Although sequence alignment of *Sm*BChE1 with the other two *Sm*ChEs revealed a single amino acid substitution in the peripheral anionic site (Glu – Trp), acyl binding pocket (Val – Leu) and catalytic triad (His – Gln), it was unclear whether these changes alone were enough to classify *Sm*BChE1 as a BChE; based on the significant difference in substrate preference, however, this classification would appear valid. Cloning of a recombinant BChE from *S. mansoni* is consistent with our observations of BChE activity in parasite extracts, and *S. mansoni* schistosomula exhibited significantly more activity than adults, as did *S. mansoni* compared to *S. haematobium* adults. It has been reported that *S. mansoni* is more sensitive to the BChE inhibitor, iso-OMPA, than *S. haematobium* [39] and it may be due to the increased BChE activity in *S. mansoni*. Indeed, this relationship has been documented between AChE and metrifonate (precursor of DDVP used in this study); *S. haematobium* is more sensitive to the inhibitor than *S. mansoni* because of the greater amount of AChE on the worm’s surface [39].

For the first time, we document the presence of secreted *Sm*ChE activity in schistosomes and AChE activity was highest in cercarial ES products. Of the intra-mammalian stages tested, AChE activity was highest in schistosomula and adults and may be acting to bind and neutralize exogenous AChE inhibitors [40] (thus protecting tegumental and somatic AChE) or host-derived ACh to mitigate the immunomodulatory effects of this molecule. Extending this hypothesis, ES products from cultured female worms had lower AChE activity than males and could be due to females worms having less of a requirement for this defensive mechanism as they reside in the relative shelter of the gynecophoric canal. BChE activity was present in the ES products of adults, schistosomula and cercariae and was significantly higher in the intra-mammalian larval stage than the other two stages. The *Sm*ChE molecules present in ES were isolated by purification on edrophonium (a reversible ChE inhibitor) sepharose and, consistent with the class of activity observed in ES, identified by mass spectrometry as *Sm*AChE1 and *Sm*BChE1; the former being forty-fold more abundant than the latter.

RNAi-mediated silencing of *smache1* and *smache3* in schistosomula showed decreases in AChE activity, consistent with reductions in transcript and protein expression levels. Moreover, inhibition of this biochemical activity was greater in schistosomula treated with the *smche* siRNA cocktail than parasites receiving any of the individual treatments, further evidence suggestive of simultaneous silencing of all *smache* paralogs. AChE activity inhibition in *smache3*-silenced parasites was more pronounced than in *smache1*-silenced parasites, which was inconsistent with protein level reductions and this may be due to the increased AChE activity reported for *Sm*AChE1 (Vmax = 5.57 nmol/min/mg, Km = 5.83 mM) compared to *Sm*AChE3 (Vmax = 5.59 nmol/min/mg, Km = 10.87 mM). It is also possible that there may not be a direct correlation between AChE activity and protein expression, given that additional, non-cholinergic functions have been ascribed to ChEs [5, 22, 41]. This may also be the reason why no significant decrease in BChE activity was observed in *smbche1*-silenced parasites, despite significant reductions in transcript and protein expression levels.

Previous studies have documented the involvement of *Sm*ChEs in the uptake of exogenous glucose by schistosomes through the ablation of the glucose uptake pathway by organophosphorus [39] and large molecule [23] AChE inhibitors, so we sought to identify the *Sm*ChE paralog(s) responsible for this mediation through the use of RNAi targeting *smche* genes. Individual gene knockdown of *smache1* and *smache3* suppressed glucose uptake in schistosomula, implying that both genes were involved in regulation of this mechanism. Tegumental AChE is speculated to mediate glucose uptake by limiting the interaction of ACh with tegumental nicotinic ACh receptors which is thought to decrease the amount of glucose uptake through surface glucose transporters. The fact that both molecules are localized to the tegument and can hydrolyze ACh therefore provides evidence for their role in this pathway. Silencing of *smbche1* in schistosomula did not show any difference in glucose uptake and is probably reflective of the molecule’s limited role in ACh hydrolysis. Transcript levels of *sgtp1* and *sgtp4* were not significantly changed in *smache1*- and *smache3*-silenced parasites, suggesting that *Sm*AChEs may facilitate glucose uptake in a manner which does not directly involve glucose transporters. Indeed, at least in nematodes, AChEs have been proposed to be involved in altering the permeability of surrounding host cells, allowing nutrients (such as glucose) to leak into the parasite niche and be uptaken [42].

Individual *smche* silencing in schistosomula resulted in significant decreases in parasite viability at various timepoints after treatment, with *smache3*-silenced parasites showing the most rapid and significant decrease in viability after treatment. Of the *smche* paralogs studied, *smche3* is the only one whose expression is significantly upregulated between *S. mansoni* cercariae and schistosomula [36], an observation consistent with qPCR data, and so silencing this relatively highly expressed gene may have the most profound effects of all *smche* silencing on parasite viability. The viability of parasites treated with all three *smche* siRNAs was significantly decreased compared to parasites treated with an individual siRNA, suggesting functional overlap exists between the paralogs. This redundancy has been documented in AChE-knockout mice where BChE has the ability to hydrolyze ACh in the absence of AChE [43, 44]. Moreover, AChE deletion is found to be lethal in *Drosophila* only because there is no alternative BChE paralog to compensate for the lack of ACh hydrolysis [45, 46]. Similar to the observations in this study, simultaneous knockdown of multiple ChE genes has been reported to have deleterious effects on their target organisms including the insects *Plutella xylostella* [47], *Chilo suppressalis* [48] and *Tribolium castaneum* [49] and the nematodes *Nippostrongylus brasiliensis* [50] and *Caenorhabditis elegans* [51]. Similarly, chemotherapy with “broad spectrum” ChE inhibitors has shown to be effective against a range of organisms, including pest insects [52], schistosomes [23, 53–55] and parasitic nematodes [56]. It is likely that the simultaneous silencing of the *smche* genes in this study has a profound effect on parasite viability due to the knockdown of cholinergic signaling, a process to which all paralogs contribute, as they have all been shown to hydrolyze ChE substrates. Also possible is that knockdown of these three genes might have resulted in the ablation of multiple other functions that have been suggested for these molecules [5, 49, 57]. Reflective of *in vitro* silencing experiments, worm recovery from mice infected with all groups of *smche*-silenced parasites was significantly less than controls, indicating that suppression of *smache1, smbche1* or *smache3* could inhibit schistosome establishment and/or development in the host. An immunomodulatory function that results in a host environment favorable to parasite survival has been suggested for AChE secreted by *N. brasiliensis* [31] and so it may be that, if schistosome ChEs have similar non-neuronal roles, impairment of ChE function in these parasites may lead to more efficient immune-mediated worm expulsion. There is now a growing body of evidence that AChEs play non-classical roles as adhesion molecules [4, 58, 59] due to these two protein families sharing significant domain homology and so *Sm*ChE-silenced worms may be unable to properly establish in their site of predilection due to impaired adherence to host vasculature. Indeed, *in vivo* treatment with AChE-inhibitory tris (p-aminophenyl) carbonium salts [60] results in a shift of the worms from the mesenteries to the liver, which may be a consequence of improper parasite attachment. Schistosomula silenced for all three *smche* genes exhibited the highest mortality *in vivo* when worm recovery was averaged across the two independent trials, suggesting that, not only is simultaneous knockdown of *Sm*ChEs required to overcome any functional redundancy between the molecules, but that this treatment has the largest impact on parasite pathogenesis due to the inhibition of multiple biological functions collectively orchestrated by these proteins.

The *smche* transcript levels of silenced worms recovered from mice were no different from control parasites and it is likely that the surviving worms received less siRNA than those worms that died in the host. Alternatively, surviving worms might have recovered from the transient effects of RNAi, highlighting the advantage of targeted gene knockout techniques such as CRISPR/Cas9 which has recently been reported for the first time in schistosomes [61].

It is generally accepted that vertebrate BChE has a predominant role in the detoxification of ingested or inhaled drugs and poisons such as the AChE-inhibitory organophosphorous esters that constitute nerve agents and pesticides due to the binding of the enzyme to these molecules (reviewed in [3]). Inactivation of *Sm*BChE1 in parasite extracts and live schistosomula by the BChE inhibitor iso-OMPA or through RNAi-mediated silencing potentiated the effects of DDVP whereas addition of exogenous *Sm*BChE1 mitigated the effects, suggesting a similar detoxification role exists for schistosome BChE as for the vertebrate enzyme. Numerous plant species in the *Solanaceae* used for their nutritional value (potatoes, for example) and others employed in traditional medicine for their anthelmintic properties contain naturally-occurring AChE-inhibitory compounds [62, 63], and so it may be that the evolution of this dietary behavior in schistosomiasis endemic populations has resulted in selective pressure on the parasite to produce these particular ChE molecules. The localization of *Sm*BChE1 to the tegument and its presence in ES products may further support this hypothesis, as the enzyme would be spatially available to interact with toxins present in the host environment, thus safeguarding parasite AChE against AChE inhibitors. Moreover, BChE activity is higher in *S. mansoni* than *S. haematobium*, which is more sensitive to the effects of the organophosphorous AChE inhibitor metrifonate. It has been reported that this sensitivity is due to the larger amount of tegumental AChE present in *S. haematobium* [39], but it may also be due to the reduced amount of BChE available to detoxify the inhibitor, as a similar relationship has been reported in studies which use human BChE to counter organophosphate toxicity [64]. Plasma-derived human BChE is currently in a phase I clinical trial as a nerve agent detoxifier, and a recombinant human BChE mutant is being used to prevent relapse in cocaine addicts due to the enzyme’s ability to hydrolyze the drug into inactive by-products [3]. One of the major limitations of these approaches, however, is the catalytic turnover of human BChE [3] and so there is emphasis on the identification of BChE homologues from other organisms, such as *Sm*BChE1, that might offer improved detoxification activity in this regard.

Inhibition of BChE in the absence of DDVP results in parasite death so a bio-scavenging role is possibly not the only function of this enzyme. Indeed, vertebrate BChE has also been shown to have roles in (1) ACh hydrolysis in situations of AChE deficiency [65], (2) fat metabolism by hydrolyzing the feeding stimulant peptide octanoyl ghrelin [66], and (3) scavenging polyproline-rich peptides to regulate protein-protein and protein-DNA interactions [67].

In summary, the work herein has identified multiple ChE paralogs in the genome of *S. mansoni* where previous studies, making use of the technology available at the time, attributed ChE activity to a single AChE. Consistent with previous observations that ChEs are multi-faceted enzymes, we posit that the three ChE paralogs described herein may fulfil distinct neuronal and non-neuronal functions based on their anatomical and temporal expression in the parasite and its ES products and the enzymatic activity of recombinant molecules. In addition to providing valuable insight into the functionality of individual ChE molecules, the study herein documents the essentiality of these proteins, providing a compelling evidence base for their use as intervention targets against schistosomiasis.

## Materials and Methods

### Ethics statement

All experimental procedures reported in the study were approved by the James Cook University (JCU) animal ethics committee (ethics approval numbers A2271 and A2391). Mice were maintained in cages in the university’s quarantine facility (Q2152) for the duration of the experiments. The study protocols were in accordance with the 2007 Australian Code of Practice for the Care and Use of Animals for Scientific Purposes and the 2001 Queensland Animal Care and Protection Act.

### Parasite maintenance, culture and ES collection

*Biomphalaria glabrata* snails infected with *S. mansoni* (NMRI strain) were obtained from the Biomedical Research Institute (BRI), MD, USA. Cercariae were shed from infected snails through exposure to light at 28°C for 1.5 h and were mechanically transformed into schistosomula [68]. To obtain adult worms, 6-8 week old male BALB/c mice (Animal Resource Centre, WA, Australia) were infected with 180 cercariae via tail penetration and adults were harvested by vascular perfusion at 7-8 weeks post-infection [69]. Both adult worms and schistosomula were cultured (10 adult pairs/ml and 2000 schistosomula/ml) at 37°C and 5% CO_2_ in serum-free modified Basch medium [70] supplemented with 4 × antibiotic/antimycotic (AA - 200 units/ml penicillin, 200 μg/ml streptomycin and 0.5 μg/ml amphotericin B) (SFB) in 6 well plates. Media containing ES products was initially collected after 3 h for schistosomula or 24 h for adults, and replenished daily thereafter. ES products were stored at −80°C. Media was thawed when needed, concentrated through Amicon centrifugation filters (Sigma) with a 3 kDa molecular weight cutoff (MWCO), buffer exchanged into phosphate buffered saline, pH 7.4 (PBS) and aliquoted. Protein concentration of ES products was determined using the Pierce BCA™ Protein Assay kit (Thermofisher). To collect cercarial ES products, freshly-shed cercariae were incubated in H_2_O (4000/ml) at 25°C for 3 h. H_2_O was filtered through Whatman filter paper (11 μm) to remove cercariae and associated debris, and ES products were concentrated, quantified and stored as described for adult and schistosomula ES products.

### Parasite extract preparation

To make PBS-soluble extracts, worms were homogenized in PBS (50 μl/adult worm pair or 50 μl/1000 schistosomula) at 4°C using a TissueLyser II (Qiagen), homogenates were incubated overnight with mixing at 4°C and the supernatants collected by centrifugation at 15,000 *g* for 1 h at 4°C. Triton X-100-soluble extracts were made from the PBS-insoluble pellets by resuspension in 1% Triton X-100, 40 mM Tris-HCl, pH 7.4, mixing overnight at 4°C and the supernatant collected by centrifugation at 15,000 *g* for 1 h at 4°C. Tegument extraction was achieved using a combination of freeze/thaw/vortex [71]. In brief, parasites were slowly thawed on ice, washed in TBS (10 mM Tris/HCl, 0.84% NaCl, pH 7.4) and incubated for 5 min on ice in 10 mM Tris/HCl, pH 7.4 followed by vortexing (5 × 1 s bursts). Subsequently, the tegumental extract was pelleted at 1000 *g* for 30 min and solubilized (3×) in 200 μl of 0.1% (w/v) SDS, 1% (v/v) Triton X-100 in 40 mM Tris, pH 7.4 with pelleting at 15,000 *g* between each wash. Protein concentration was determined using the Pierce BCA Protein Assay kit, aliquoted and stored at −80°C until use.

### Bioinformatics

Based on Pfam analysis (search = cholinesterase) of the *S. mansoni* genome (http://www.geneDB.org/Homepage/Smansoni), three *smche* paralogs (*smache1* - Smp_154600, *smbche1* - Smp_125350 and *smache3* - Smp_136690) were identified. Homologous ChE sequences from other species were identified using BLASTP. (http://blast.ncbi.nlm.nih.gov/Blast.cgi) and the resulting sequences were used to generate a multiple sequence alignment using Clustal Omega (https://www.ebi.ac.uk/Tools/msa/clustalo/). MEGA 7 was used to generate a neighbor-joining tree with the Poisson correction distance method and a bootstrap test of 1,000 replicates [72]. The tree was visualized with The Interactive Tree of Life (iTOF) online phylogeny tool (https://itol.embl.de/). Structure-homology 3D models of *Sm*ChEs were generated using the I-TASSER server (http://zhanglab.ccmb.med.umich.edu/I-TASSER/). For structure visualization and catalytic triad analyses, the Accelrys Discovery Studio (Accelerys Inc.) and UCSF Chimera MatchMaker ver. 1.4 (University of California) software packages were utilized.

### Real-time qPCR

Real-time qPCR was used to assess developmental expression of *smche* genes and to determine *smche* transcript suppression resulting from RNAi experiments. RNA from miracidia, sporocysts, cercariae, adult male worms, adult female worms, and eggs were obtained from BRI. Schistosomula were cultured as described above, harvested (1,000 parasites) after either 3 h, 24 h, 3 or 5 days, washed three times in PBS and stored at −80°C until use. Schistosomula from RNAi experiments were similarly processed. Total RNA extraction was performed using the Trizol (Thermofisher) reagent according to manufacturer’s instructions. After air-drying, RNA pellets were re-suspended in 12 μl diethylpyrocarbonate (DEPC)-treated water. Concentration and purity of RNA was determined using an ND2000 Nanodrop spectrophotometer (Thermofisher). Synthesis of cDNA was carried out with 1 μg of total RNA using Superscript-III-Reverse Transcriptase (Invitrogen) according to the manufacturer’s instructions. Finally, cDNA was quantified, diluted to 50 ng/μl, aliquoted and stored at −20°C. Real-time qPCR primers for each *smche* (supplementary Table 1) were designed using Primer3 (http://frodo.wi.mit.edu/). The housekeeping gene *smcox1* was selected as an internal control to normalize relative *smche* gene expression [73]. Each qPCR (1 μl (50 ng) of cDNA, 5 μl of 2x SYBR green master mix (Bioline), 1μl (5 pmol/μl) each of forward and reverse primers and 2 μl of nuclease-free water) was run in a Rotor-Gene Q thermal cycler (Qiagen) using 40 cycles of 95°C for 10 seconds, 50-55°C for 15 seconds and 72°C for 20 seconds. Stage-specific *smche* gene expression levels were normalized against *smcox1* gene expression using the comparative 2^-ΔΔCT^ method [74]. All results represent the average of 5 independent experiments with data presented as mean ± SEM.

### Cloning, expression and purification of smche gene fragments in E. coli

Complete ORFs for *smache1, smbche1* and *smache3* were synthesized by Genewiz. Attempts to express full-length sequences in *E. coli* were unsuccessful, so primer sets incorporating *Nde*I (forward primer) and *Xho*I restriction enzyme sites (reverse primer) were designed (supplementary Table 1) to amplify partial, non-conserved regions of each *smche*, which might prove more amenable to expression. Sequences (containing *Nde*I/*Xho*I sites) for each p*Sm*ChE were amplified from each full-length template by PCR and cloned into the pET41a expression vector (Novagen) such that the N-terminal GST tag was removed. Protein expression was induced for 24 h in *E. coli* BL21(DE3) by addition of 1 mM Isopropyl beta-D-1-thiogalactopyranoside (IPTG) using standard methods. Cultures were harvested by centrifugation (8,000 *g* for 20 min at 4°C), re-suspended in 50 ml lysis buffer (50 mM sodium phosphate, pH 8.0, 300 mM NaCl, 40 mM imidazole) and stored at −80°C. Cell pellets were lysed by three freeze-thaw cycles at −80°C and 42°C followed by sonication on ice (10 × 5 s pulses [70% amplitude] with 30 s rest periods between each pulse) with a Qsonica Sonicator. Triton X-100 was added to each lysate at a final concentration of 3% and incubated for 1 h at 4°C with end-over-end mixing. Insoluble material (containing r*Sm*ChEs) was pelleted by centrifugation at 20,000 *g* for 20 min at 4°C. The supernatant was discarded, and inclusion bodies (IBs) were washed twice by resuspension in 30 ml of lysis buffer followed by centrifugation at 20,000 *g* for 20 min at 4°C. IBs were then solubilized sequentially by resuspension in 25 ml lysis buffers containing either 2 M, 4 M or 8 M urea, end-over-end mixing overnight at 4°C and centrifugation at 20,000 *g* for 20 min at 4°C. Finally, supernatant containing solubilized IBs was diluted 1:4 in lysis buffer containing 8M urea and filtered through a 0.22 μm membrane (Millipore). Solubilized IBs were purified by immobilized metal affinity chromatography (IMAC) by loading onto a prepacked 1 ml His-Trap HP column (GE Healthcare) equilibrated with lysis buffer containing 8M urea at a flow rate of 1 ml/min using an AKTA-pure-25 FPLC (GE Healthcare). After washing with 20 ml lysis buffer containing 8M urea, bound His-tagged proteins were eluted using the same buffer with a stepwise gradient of 50-250 mM imidazole (50 mM steps). Fractions containing r*Sm*ChEs (as determined by SDS-PAGE) were pooled and concentrated using Amicon Ultra-15 centrifugal devices with a 3 kDa MWCO and quantified using the Pierce BCA Protein Assay kit. The final concentration of each r*Sm*ChE was adjusted to 1 mg/ml and proteins were aliquoted and stored at −80°C.

### Generation of anti-rSmChE antisera and purification of IgG

Three groups of five male BALB/c mice (6-week-old) were intraperitoneally immunized with either r*Sm*AChE1, r*Sm*BChE1 or r*Sm*AChE3 subunits (50 μg/mouse). Antigens were mixed with an equal volume of Imject alum adjuvant (Thermofisher) and administered three times, two weeks apart. Two weeks after the final immunization, mice were sacrificed and blood was collected via cardiac puncture. Blood from all mice in each group was pooled and serum was separated by centrifugation after clotting and stored at −20°C. Polyclonal antibodies were purified from mouse sera using Protein A Sepharose-4B (Thermofisher) according to the manufacturer’s instructions. Serum from naïve mice was similarly processed.

### Immunolocalization using anti-rSmChE antisera

#### Adult worm sections

Freshly perfused adult *S. mansoni* and *S. haematobium* worms were fixed in 4% paraformaldehyde, embedded in paraffin and sections (7 μm thick) were cut in a cryostat. Following deparaffinization in xylene and rehydration in an ethanol series, antigen retrieval was performed by boiling the slides in 10 mM sodium citrate, pH 6.0, for 40 min followed by a solution of 10 mM Tris, 1 mM EDTA, 0.05% Tween, pH 9.0, for 20 min. All sections were then blocked with 10% heat-inactivated goat serum for 1 h RT. After washing 3 times with PBST, sections were incubated with anti-*Sm*AChE1, anti-*Sm*BChE1, anti-*Sm*AChE3, naïve sera (negative control), *S. mansoni* or *S. haematobium* infected mouse sera (positive controls) (1:50 in PBST) overnight at 4°C and then washed again (3 × 5 min each). Finally, the sections were incubated with goat-anti-mouse IgG-alexafluor647 (Sigma) (1:200 in PBST) for 1 h in the dark at RT. After a final washing step, slides were mounted with coverslips in Entellan mounting medium (Millipore). Fluorescence and bright-field microscopy were performed with an AxioImager M1 fluorescence microscope (Zeiss) using 10× and 20× objectives.

#### Live schistosomula

*In vitro* cultured living cercariae and schistosomula (3 h, 24 h, 3 and 5 days old) were harvested, washed with PBS and then blocked with PBST/10% heat-inactivated goat serum for 30 min at RT. Following three washes, the larvae were incubated with anti-*Sm*AChE1, anti-*Sm*BChE1, anti-*Sm*AChE3 or naïve serum (negative control) (1:100 in PBST) overnight at 4°C. Parasites were washed again before incubation with goat-anti-mouse IgG-alexafluor647 (Sigma) (1: 200 in PBST) for 1 h in the dark at RT, followed by 3 washes. Finally, schistosomes were fixed in 4% paraformaldehyde and transferred to a microscope slide for fluorescence microscopy using an AxioImager M1 fluorescence microscope.

### Cloning and expression of full-length SmChEs in P. pastoris

Full-length sequences (minus the signal peptide) of *Sm*AChE1, *Sm*BChE1 and *Sm*AChE3 (f*Sm*ChEs) were *Eco*RI/*Xba*I cloned into the C-terminal 6-His-tagged pPICZαA expression vector (Invitrogen) to facilitate secretory expression. Recombinant plasmids (20 μg) were linearized with Pmel (f*Sm*AChE1 & f*Sm*BChE1) and SacI (f*Sm*AChE3), purified by ethanol precipitation and resuspended in 15 μl of H_2_O. Linearized vectors were electroporated according to the manufacturer’s instructions into *P. pastoris* X-33 cells (Thermofisher) in 2 mm cuvettes (2 ms, 2000V, 25 μF, 200 Ω, square wave pulse), using a Gene Pulser Xcell (Bio-Rad), plated onto YPDS agar plates containing 100 μg/ml zeocin and incubated for 3 days at 30°C. Resultant colonies were then picked and patched onto YPDS agar containing 2 mg/ml zeocin and plates incubated at 30°C until colonies were visible. A high-expressing clone of each r*Sm*ChE (determined from pilot expression experiments) was used to inoculate 5 ml BMGY media supplemented with 50 ug/ml zeocin and grown overnight at 30°C with rotation at 250 rpm. The entire culture was then used to inoculate 250 ml of BMGY in a 2L baffled flask and incubation was continued for 24 h at 30°C. Cells were pelleted at 5000 *g* for 20 min at RT, re-suspended in 1L of BMMY (to induce protein expression) and split between 2 × 2L baffled flasks, which were incubated with shaking (250 rpm) at 30°C for 72 h. Methanol was added to a final concentration of 0.5% (2.5 ml/flask) every 24 h to maintain induction of protein expression. Culture medium containing the secreted f*Sm*ChE proteins was harvested by centrifugation (5000 *g* for 20 min at RT) and filtered through a 0.22 μm membrane filter (Millipore). Recombinant proteins were purified by IMAC using an AKTA-pure-25 FPLC (GE Healthcare). Briefly, culture medium was loaded onto a 5 ml His-excel column, pre-equilibrated with binding buffer (50 mM PBS pH 7.4, 300 mM NaCl), washed with 20 column volumes of binding buffer and then eluted with binding buffer containing a linear imidazole gradient (20 to 500 mM). The purity of fractions within the main peak was analyzed by SDS-PAGE and fractions of appropriate purity were pooled, concentrated and buffer exchanged into PBS using Amicon Ultra-15 centrifugal devices with a 3 kDa MWCO and quantified using the Pierce BCA Protein Assay kit. The final concentration of each f*Sm*ChE was adjusted to 1 mg/ml and proteins were aliquoted and stored at −80°C.

### SmChE enzyme assays

Activity of f*Sm*ChEs, extracts and ES samples was determined by the Ellman method [75]; modified for use with 96 well microplates. Samples (parasite extracts, ES and f*Sm*ChEs) were diluted in assay buffer (0.1M sodium phosphate, pH 7.4), and 2 mM acetylthiocholine (AcSCh) or butyrylthiocholine (BcSCh) (Sigma) and 0.5 mM 5, 5’-dithio-bis (2-nitrobenzoic acid) (DTNB) (Sigma) was added. The absorbance increase was monitored every 5 min at 405 nm in a Polarstar Omega microplate reader (BMG Labtech). Specific activity was calculated using the initial velocity of the reaction and extinction coefficient of 13,260 M^−1^ cm^−1^ for TNB. To investigate sensitivity of parasite ES products to AChE inhibitors, 25 μg of adult ES was pre-treated with 1 μM DDVP - active metabolite of the organophosphorous AChE inhibitor metrifonate - for 20 min at RT before measuring activity. Kinetic parameters of f*Sm*ChEs were characterized by measuring enzyme activity at differing substrate concentrations and plotting enzyme activity [V] vs. substrate concentration [S]. The Km ([S] at 1/2 Vmax) was calculated using the Michaelis Menton equation. Enzyme assays with inhibitors were performed as above except that f*Sm*ChEs in assay buffer were pre-treated with 1 μM DDVP, in the case of f*Sm*AChE1 and f*Sm*AChE3, or 1 mM iso-OMPA – a membrane-impermeable specific BChE inhibitor - in the case of f*Sm*BChE1, for another 20 min at RT. Experiments were performed in triplicate with data presented as the mean ± SEM.

### Purification of secreted SmChEs from adult S. mansoni ES products

Affinity chromatography using edrophonium chloride-sepharose was used to purify *Sm*AChE from *S. mansoni* based on the method of Hodgson and Chubb [76]. Briefly, 1 *g* of epoxy-activated sepharose 6B beads was washed with distilled H_2_O, the slurry centrifuged at 814 *g* for 5 min and the pellet gently resuspended in 50 mM sodium phosphate, pH 8.0, containing 200 mM edrophonium chloride (1:2 ratio of sepharose:edrophonium chloride). The pH of the solution was adjusted to 10.0 and coupling of edrophonium with the sepharose was facilitated by incubating the mixture overnight with shaking at 50°C. The gel was then washed sequentially with 10 volumes each of 100 mM sodium acetate, pH 4.5, 12 mM sodium borate, pH 10.0, and distilled H_2_O and finally resuspended in distilled H_2_O to generate a 1 ml gel slurry. The gel slurry was packed into a chromatography column (10 cm long, 1 cm diameter) and equilibrated by gravity flow at 4°C with 10 column volumes (CV) of equilibration buffer (50 mM phosphate buffer, pH 8.0). Approximately 20 ml of ES from adult *S. mansoni* (concentrated through a 10 kDa MWCO centrifugal filter from a starting volume of 500 ml of media, harvested each day for 7 days from 100 pairs of adult worms and buffer exchanged into equilibration buffer) was added to the column followed by washes with 20 CV of equilibration buffer and 20 CV of equilibration buffer containing 500 mM NaCl. Bound *Sm*ChE was then eluted with 10 CV of equilibration buffer containing 500 mM NaCl and 20 mM edrophonium chloride. The eluate was concentrated and buffer exchanged into PBS using a 10 kDa MWCO centrifugal filter (edrophonium chloride is an AChE inhibitor and would interfere with subsequent activity assays) and resolved by 10% SDS-PAGE to check purity and facilitate identification by mass spectrometry.

### Mass spectrometric analysis of purified, secreted SmChE

Bands of interest were manually excised from the SDS polyacrylamide gel, washed with 50% acetonitrile and dried under vacuum at 30°C. Cysteine residues were reduced with 20 mM DTT for 1 h at 65°C followed by alkylation with 50 mM iodoacetamide for 40 min at 37°C in the dark. In-gel trypsin digestion was performed at 37°C overnight with 0.8 ng of trypsin in trypsin reaction buffer (40 mM ammonium bicarbonate, 9% acetonitrile). The supernatant was removed to a fresh microfuge tube and stored at 4°C, and the remaining peptides were further extracted from the gel pieces by incubation with 0.1% trifluoracetic acid (TFA) at 37°C for 45 min. The newly extracted supernatant was combined with the previously collected supernatant, then dried under vacuum. Prior to the matrix-assisted laser desorption/ionization-time-of-flight mass spectrometry (MALDI-TOF MS) analysis, peptides were concentrated and desalted using ZipTips (Millipore) following the manufacturer’s instructions. Tryptic peptides were re-dissolved in 10 μl 5% formic acid and 6 μl was injected onto a 50 mm 300 μm C18 trap column (Agilent Technologies) followed by an initial wash step with Buffer A (5% (v/v) ACN, 0.1% (v/v) formic acid) for 5 min at 30 μl/min. Peptides were eluted at a flow rate of 0.3 μl/min onto an analytical nano HPLC column (15 cm × 75 μm 300SBC18, 3.5 μm, Agilent Technologies). The eluted peptides were then separated by a 55-min gradient of buffer B (90/10 acetonitrile/ 0.1% formic acid) 1-40% followed by a 5 min steeper gradient from 40-80%. The mass spectrometer (ABSciex 5600 Triple Tof) was operated in data-dependent acquisition mode, in which full scan TOF-MS data was acquired over the range of 350-1400 m/z, and over the range of 80-1400 m/z for product-ion observed in the TOF-MS scan exceeding a threshold of 100 counts and a charge state of +2 to +5. Analyst 1.6.1 (ABSCIEX) software was used for data acquisition and analysis.

For protein identification, a database was built using the *S. mansoni* genome v5.0 [http://www.genedb.org/Homepage/Smansoni] with the common repository of adventitious proteins (cRAP, http://www.thegpm.org/crap/) appended to it. Mascot v.2.5.1 (Matrix Science) was used for database search. Carbamidomethylation of Cys was set as a fixed modification and oxidation of Met and deamidation of Asn and Gln were set as variable modifications. MS and MS/MS tolerance were set at 10 ppm and 0.1 Da, respectively and only proteins with at least two unique peptides (each composed of at least seven amino acid residues) identified were considered reliably identified and used for analysis.

### siRNA design and synthesis

Three short interfering RNA duplexes (siRNAs) targeting each of the three identified *smche* paralogs were designed (supplementary Table 2) and checked to avoid off-target silencing by BLAST search using the *S. mansoni* genome. An irrelevant siRNA from firefly luciferase (*luc*) was selected as a negative control [77]. All siRNAs were commercially synthesized (Integrated DNA Technologies) and oligonucleoitdes were suspended to a concentration of 1 μg/μl in DEPC-treated water.

### Electroporation of schistosomula with siRNA

Prior to electroporation, mechanically transformed schistosomula were cultured for 24 h (2,000 schistosomula/ml), at 37°C and 5% CO_2_ in SFB in 6 well plates. After 3 washes with PBS, schistosomula were re-suspended in modified Basch medium (3,000 schistosomula/100 μl) and 3,000 schistosomula were transferred into a Genepulser 4 mm electroporation cuvette (Bio-Rad) for every siRNA treatment (four) and timepoint (four for each siRNA treatment – 1, 3, 5 and 7 days). Schistosomula were electroporated with 10 μg of either *luc, smache1, smbche1* or *smache3* siRNA or a combination of all three *smche* siRNAs (30 μg total) using a Bio-Rad Gene Pulser Xcell (single 20 ms pulse – 125 V, 25 μF capacitance, 200 Ω resistance, square wave electroporation) at RT, added to 24 well plates containing 1 ml pre-warmed SFB and incubated (37°C, 5% CO_2_) for 7 days. Schistosomula were harvested at each timepoint and approximately 1,000 parasites were used for qPCR analysis (to assess transcript knockdown), 1,700 parasites were used for protein extract preparation (to examine phenotypic knockdown) and 300 parasites were used for Trypan Blue exclusion assays (to determine parasite viability). All parasite material was generated and separately analyzed from 2 independent experiments.

### Determination of viability in siRNA-treated schistosomula

Schistosomula (100 parasites/replicate) were harvested at each timepoint and viability was determined by Trypan Blue exclusion staining [78]. Briefly, schistosomula were stained with 0.16% Trypan Blue in PBS with gentle shaking for 30 min at RT and then excess stain was removed by multiple washes in PBS before fixing in 10% formalin. Parasites were counted under 10× objective and live parasites (which had not taken up stain) were expressed as a percentage of total worms. Each assay was performed in triplicate.

### Evaluation of protein expression in siRNA-treated schistosomula

Western blots were performed with day-7 parasite extracts (20 μg) following standard procedures. The blots were probed with the anti-*Sm*ChE antibodies (1:1000 in PBST) generated herein. A polyclonal anti-Sm-paramyosin antibody [77] was used as a loading control.

### Glucose uptake in schistosomula treated with siRNA

In a separate RNAi experiment, newly transformed schistosomula (5,000/treatment) were incubated for 5 days in SFB. Parasites were then electroporated with siRNAs as described above and finally transferred to serum-free DMEM (1 mg/ml glucose) supplemented with 4’AA. Media (50 μl) from each experiment was collected 72 h post-treatment and the amount of glucose was quantified using a colorimetric glucose assay kit (Sigma) following the manufacturer’s instructions. Parasite viability at this timepoint was determined by Trypan Blue exclusion and transcript levels of each *smche*, as well as the glucose transporters *sgtp1* and *sgtp4*, were also measured. Glucose levels were normalized according to the number of parasites and expressed relative to the *luc* group. Data is the average of 2 biological and 3 technical replicates ± SEM.

### Infection of mice with SmChE siRNA-treated schistosomula

One-day-old schistosomula (10,000) were electroporated as above in 500 μl of SFB with 50 μg of either *luc, smache1, smbche1* or *smache3* siRNA or a combination of all three *smche* siRNAs (150 μg total). Parasites were injected intramuscularly into both thighs (1,000 per thigh) of male 6-8 week BALB/c mouse (5 mice per treatment group) using a 23-gauge needle. A control group of mice were similarly injected with non-electroporated schistosomula. Adult worms were perfused 20 days later to assess the number of worms that had matured and reached the mesenteries. Experiments were performed independently in duplicate. After each experiment, transcript levels of each *smche* from surviving worms were assessed using real-time qPCR.

### Bio-scavenging of carboxylic esters by SmBChE1

To test the hypothesis that *Sm*BChE1 may play a role in the bio-scavenging of AChE-inhibitory molecules, we first sought to determine whether inhibition of BChE activity would potentiate the AChE-inhibitory and anti-schistosome effects of organophosphates (OP)s. Schistosomula extracts (20 μg) were diluted in assay buffer, then iso-OMPA was added to a final concentration of either 1 or 2 mM and incubated for 20 min at RT. DDVP was then added to a final concentration of 1 μM and the samples were further incubated for 20 min at RT; the final reaction volume was 180 μl. ACh (final concentration 2 mM) and DTNB (final concentration 0.5 mM) were then added and the absorbance was monitored every 5 min at 405 nm in a Polarstar Omega microplate reader (BMG Labtech). Extracts that were not treated with iso-OMPA with or without DDVP treatment were used as controls. Experiments were performed in triplicate with data presented as the mean ± SEM.

The same experiments were performed on live schistosomula using either an inhibitor- or RNAi-based approach. For the inhibitor-based experiment, 24 h schistosomula (1,000/treatment in 1 ml SFB) were pretreated with iso-OMPA at the non-lethal concentration of 100 μM and, 1 h after iso-OMPA treatment, schistosomula were treated with 1 μM DDVP and cultured for 5 h at 37°C in 5% CO_2_. Parasites that were not treated with iso-OMPA but treated with DDVP were used as controls. For the RNAi-based experiment, 24 h schistosomula (1,500/100 μl SFB) were electroporated with 10 μg of either *smbche1* or *luc* siRNA as described above, added to 24 well plates containing 1 ml pre-warmed SFB and incubated (37°C, 5% CO_2_) for 3 days before being treated with 1 μM DDVP and cultured for a further 5 h. For both inhibitor- and RNAi-based experiments, schistosomula viability was determined using Trypan Blue staining and data is presented as the mean ± SEM of 2 biological and 3 technical replicates.

In a reverse testing of the bio-scavenging hypothesis, we sought to determine whether addition of *Sm*BChE could mitigate the effects of DDVP. Ten micrograms of f*Sm*BChE1 was pre-incubated with 1 μM final concetration DDVP in AChE assay buffer (170 μl final volume) for 20 min at RT. Schistosomula extracts (20 μg), ACh (final concentration 2 mM) and DTNB (final concentration 0.5 mM) were then added and the absorbance was monitored every 5 min at 405 nm iichlorvosn a Polarstar Omega microplate reader. Reactions without f*Sm*BChE or without DDVP were used as controls. Experiments were performed in triplicate with data presented as the mean ± SEM. Again, the same experiments were performed on live schistosomula. After the pre-treatment of different amounts of f*Sm*BChE (10, 5, and 2.5 μg) with 1 μM DDVP in 500 μl SFB, 24 h schistosomula (1,000/treatment in 500 μl SFB) were added, incubated at 37°C and 5% CO_2_ for 24 h and then parasite viability was measured by Trypan Blue staining. Experiments where a similarly expressed and purified, but irrelevant, protein (*Sm*TSP2) was used instead of f*Sm*BChE, and schistosomula cultured in media alone, were used as controls. Data is presented as the mean ± SEM of 2 biological and 3 technical replicates.

### Statistical analyses

Data were reported as the means ± SEM. Statistical differences were assessed using the student’s *t* test. *P* values less than 0.05 were considered statistically significant.

## Supporting information

**Figure S1. Regional amino acid sequence alignment of *Sm*BChE1 and its human and other helminth homologs.** Accession numbers: *Schistosoma mansoni* (*Sm*BChE1 – Smp_125350), *Schistosoma rodhaini* (SROB_0000329201), *Schistosoma haematobium* (KGB33101), *Schistosoma japonicum* (Sjp_0015690), *Clonorchis sinensis* (csin111679), *Echinostoma caproni* (ECPE_0000670801), *Fasciola hepatica* (PIS83327.1), *Hymenolepis diminuta* (HDID_0000005301), *Echinococcus granulosus* (EGR_07475.1), *Taenia solium* (TsM_000234300), *Taenia saginata* (TSAs00071g07627m00001), *Trichuris muris* (TMUE_3000012587), *Trichuris trichiura* (TTRE_0000364501), *Trichuris suis* (M514_03850), *Nippostrongylus brasiliensis* (NBR_0000102801), *Caenorhabditis elegans* (Y48B6A.8.1). Red box = catalytic triad residue.

**Figure S2. Magnified view of 3D models showing the catalytic triads of *Sm*AChE1, *Sm*BChE1, and *Sm*AChE3.** The amino acid residues of the catalytic triad of each paralog are magnified and their position number is given according to *Torpedo* AChE numbering: *Sm*AChE1 (Ser277, Gln538, Glu406), *Sm*BChE1 (Ser244, Gln538, Glu406), and *Sm*AChE3 (Ser239, His514, Glu375).

**Figure S3. Relationship between *Sm*ChEs and other invertebrate and vertebrate species.** Evolutionary history was inferred using the Neighbor-Joining method and the phylogenetic tree was generated using a ClustalW alignment. The evolutionary distances were computed using the Poisson correction method and are in the units of the number of amino acid substitutions per site. All positions containing gaps and missing data were eliminated, making for a total of 236 positions in the final dataset. The three *Sm*ChEs are indicated by bold font inside a red box. Accession numbers: *Schistosoma mansoni* (Sm_AChE1 - Smp_154600, Sm_BChE1 - Smp_125350, Sm_AChE3 - Smp_136690); *Schistosoma bovis* (Sb_AChE1 - AAQ14323); *Schistosoma haematobium* (Sh_AChE1 - AAQ14322, Sh_AChE2 - KGB33101, Sh_AChE3 - KGB33661); *Schistosoma japonicum* (Sj_AChE1 - ANH56887, Sj_AChE2 - Sjp0045440.1); *Clonorchis sinensis* (Cs_AChE1 - GAA52478, Cs_AChE2 - GAA53463, Cs_AChE3 - GAA27255); *Opisthorchis viverrini* (Ov_AChE - XP009170845, Ov_AChE - XP009168237, Ov_AChE - XP009170760); *Echinococcus granulosus* (Eg_AChE1 - JN662938, Eg_AChE2 - EgG000732400); *Hymenolepis microstoma* (Hm_AChE1 - LK053025); *Taenia solium* (Ts_AChE1 - TsM000234300, Ts_AChE - TsM001220100, Ts_AChE - TsM000001700); *Anopheles gambiae* (Ag_AChE1 - AGM16375); *Aedes aegypti* (Ae_AChE - AAB35001); *Culex tritaeniorhynchus* (Ct_AChE - BAD06210); *Caenorhabditis elegans* (Ce_AChE1 - NP510660, Ce_AChE2 - NP491141, Ce_AChE3 - NP496963); *Trichuris muris* (Tm_AChE1 - TMUEs0033000600); *Nippostrongylus brasiliensis* (Nb_AChE1 - AAK44221, Nb_AChE2 - AAC05785, Nb_AChE3 - AAK44221); *Homo sapiens* (Hs_AChE - NP000656); *Torpedo californica* (Tc_AChE - CAA27169); *Danio rerio* (Dr_AChE - NP571921); *Mus musculus* (Mm_AChE - CAA39867); *Rattus norvegicus* (Rn_AChE - NP742006).

**Figure S4. Phylogenetic analysis of *Sm*BChE1 and its human and other helminth homologs.** The phylogenetic tree was built using the maximum likelihood method with *Sm*BChE1 and the top 16 helminth ChE homologs identified from the BLASTp search, as well as human BChE. Accession numbers: *Schistosoma mansoni* (*Sm*BChE1 – Smp_125350), *Schistosoma rodhaini* (SROB_0000329201), *Schistosoma haematobium* (KGB33101), *Schistosoma japonicum* (Sjp_0015690), *Clonorchis sinensis* (csin111679), *Echinostoma caproni* (ECPE_0000670801), *Fasciola hepatica* (PIS83327.1), *Hymenolepis diminuta* (HDID_0000005301), *Echinococcus granulosus* (EGR_07475.1), *Taenia solium* (TsM_000234300), *Taenia saginata* (TSAs00071g07627m00001), *Trichuris muris* (TMUE_3000012587), *Trichuris trichiura* (TTRE_0000364501), *Trichuris suis* (M514_03850), *Nippostrongylus brasiliensis* (NBR_0000102801), *Caenorhabditis elegans* (Y48B6A.8.1).

**Figure S5. Transcript levels of glucose transporters *sgtp1* and *sgtp4* and each *smche* in individual and cocktail *smche* siRNA-treated schistosomula.** Transcript levels of each *smche* and *sgtp* in parasites treated with *smche* siRNAs are shown relative to *smche* transcript expression in schistosomula treated with the *luc* control siRNA (dashed line) and represent the mean ± SEM of triplicate qPCR assays from 2 biological replicates of each treatment). Transcript expression in all parasites was normalized with the housekeeping gene, *smcox1*. Differences in transcript levels (relative to the *luc* control) were measured by the student’s *t* test. \**P* ≤ 0.05, \*\**P* ≤ 0.01, \*\*\**P* ≤ 0.001.

**Table S1.**
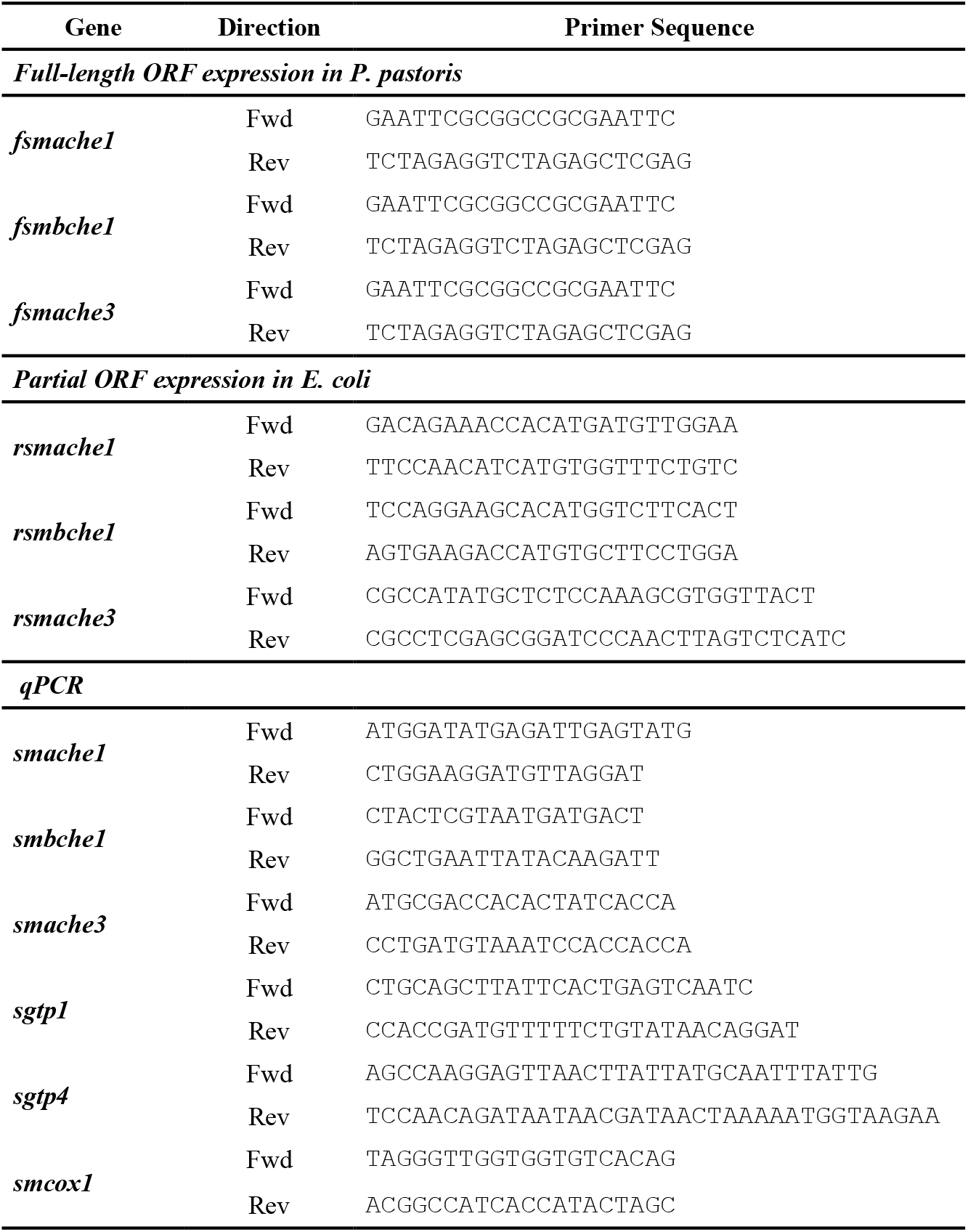
Primers used in this study

**Table S2.**
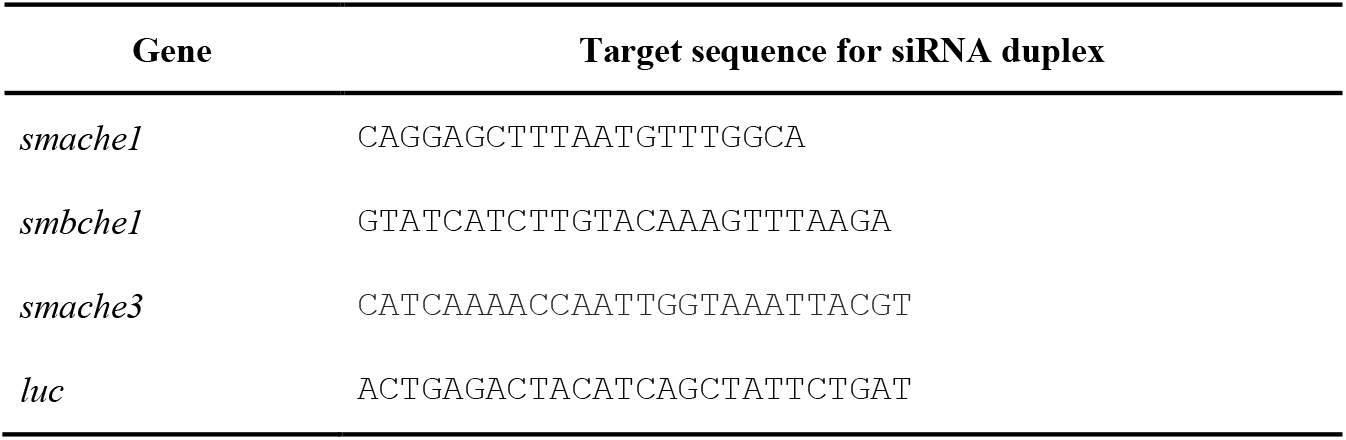
Target sequences used to design siRNA duplexes

**Table S3.**
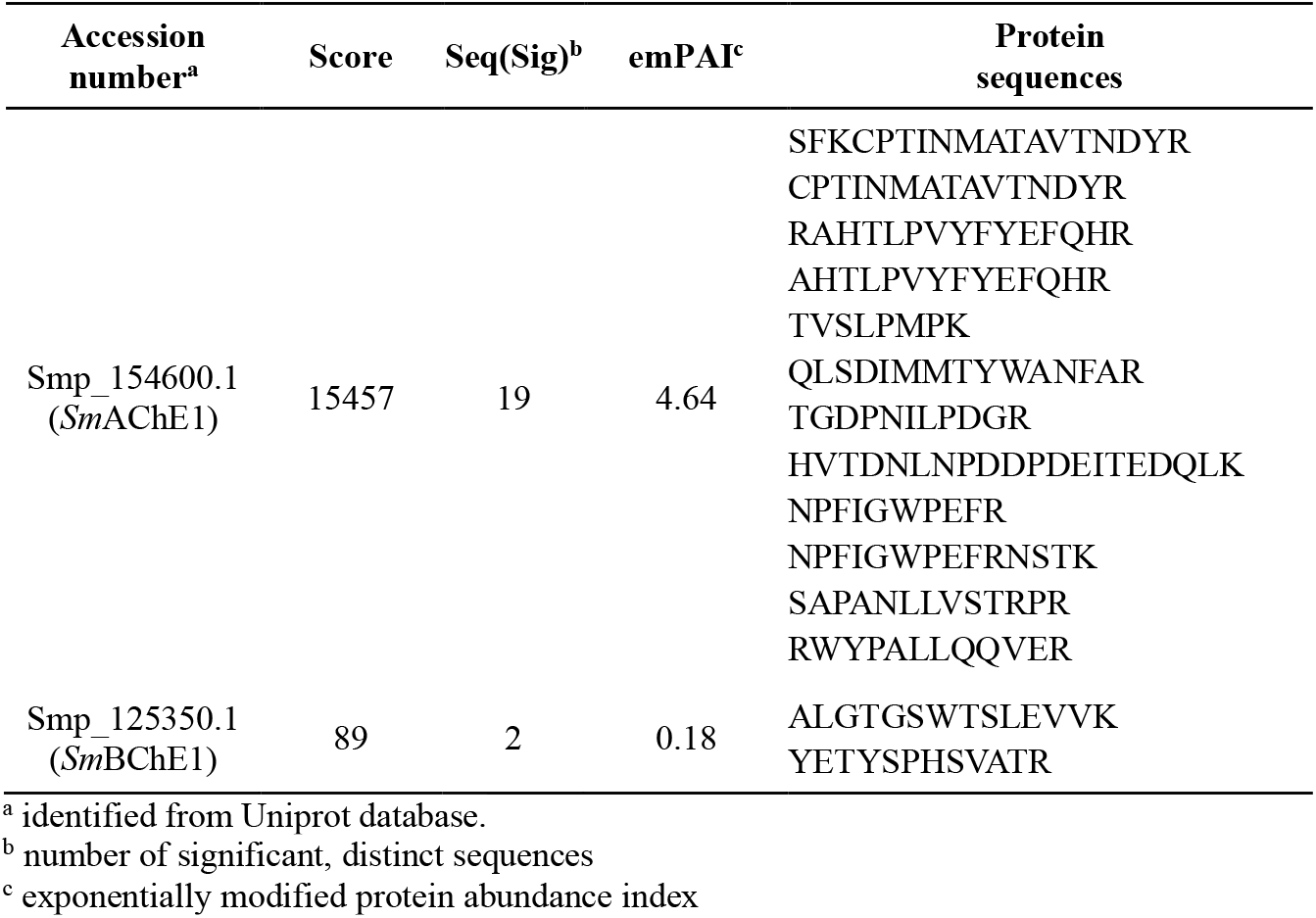
Identification by LC-MS/MS of *Sm*ChEs purified from adult *S. mansoni* ES products

