## Supplementary material for "Novel cholinesterase paralogs of *Schistosoma mansoni* have perceived roles in cholinergic signaling, glucose scavenging and drug detoxification and are essential for parasite survival": figure S1

```
H_sapiens          423 SKLPWPEWMGVM H GYEIEFVFGLPLER-----RDNYTKAEEILSRSIVKRWANFAKYGNP
C_elegans          425 SANPWPKWTGVM H GYEIEYVFGVPLHN----TTAGYTKEEMDVSEKVIDFWTTFANTGVP
N_brasiliensis     424 SANPWPKWTGVM H GYEIEYVFGVPIYN----ESAGYTKREQVLSEKIIQYWSSFEPCCTF
T_trichiura        430 SQQVWPEWMGAV H GYEINFIYGEPLNI----HRYAYTEAEKDLSRRFMSYWANFARTGNP
T_muris            430 SQQVWPEWMGAV H GYEINFVYGEPLNV----RQYAYTEAEKELSRRFMRYWANFARTG--
T_suis             430 SQQVWPEWMGAV H GYEINFIYGEPLNI----HRYAYTEAEKDLSRRFMSYWANFARTG--
Sm_BChE1           481 SGLTWPQWTGIM Q GYEAEYIFGAPFNQAFTDNYYNFTLEEKRLSEEMMQFWTNFASTGSP
S_haematobium      481 TGLTWPEWTGVM Q GYEAEYIFGAPFNQAFIDHYYNFTLEEKRLSEEIMQFWTNFASTGSP
S_japonicum        481 SGLTWPEWTGVM Q GYEAEYIFGAPFNQAFTDNYYNFTPEEKRLSEEMMQFWTNFASTG--
S_rodhaini         404 SGLTWPQWTGIM Q GYEAEYIFGAPFNQAFTDNYYNFTLEEKRLSEEMMQFWTNFASTGSP
C_sinensis         305 QASPWPQWTGVM Q GYEAEYIFGAPFNPDYQKQFYNFTDEERRLSEEMMRFWTNFASTGSP
E_caproni          336 EALSWPEWTGVM Q GFEAEYIFGAPFNPDFQQQFHNFTEDEKRLSEEMMRCWTNFASTG--
T_solium           480 SGWTWPNWTGVM Q AYEAEYIFGAPLNLKFQMDFYKFSDEERKLSASIMQYWANFAATGSP
T_saginata         481 SGWTWPNWTGVM Q AYEAEYIFGAPLNLKFQMDFYKFSDEERKLSASIMQYWANFAATGSP
E_granulosus       410 SGWTWPNWTGVM Q AYEAEYIFGAPLNLKFQMDFYKFSDEERKLSASIMQYWANFAATGSP
H_diminuta         477 SGWTWPNWTGVM Q GYEAEYVFGAPLNFKFQSEFYKFSEEERLFSNQIMQFWANFAATGSP
F_hepatica         456 VAFSWPEWTGVM Q GFEAEYIFGAPFNPDFQREFYNFTDEEKRLSEEMMRCWTNFASTGSP
```
