## Supplementary figures and images for "Novel cholinesterase paralogs of *Schistosoma mansoni* have perceived roles in cholinergic signaling, glucose scavenging and drug detoxification and are essential for parasite survival"

### figure S2

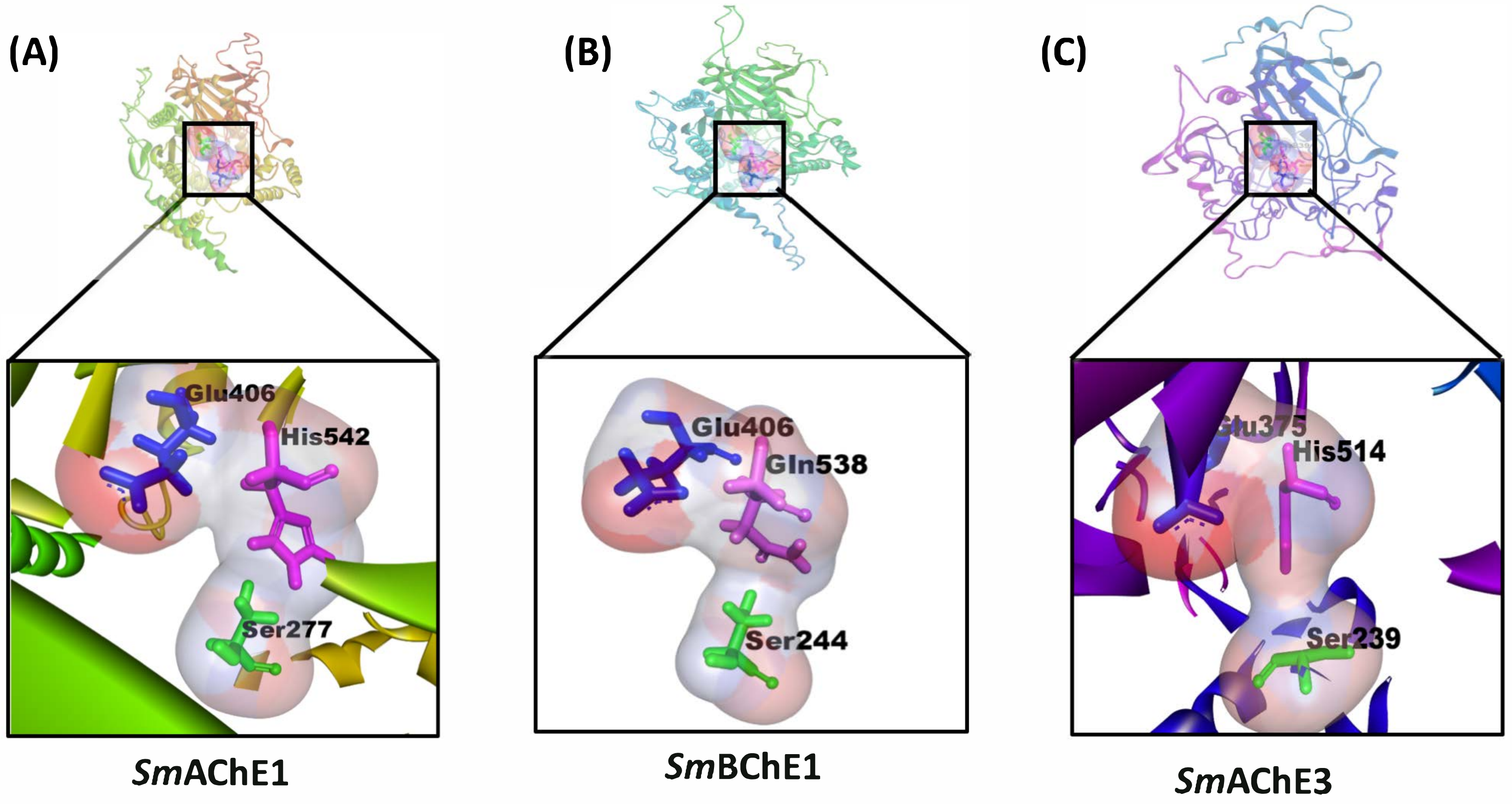

### figure S3

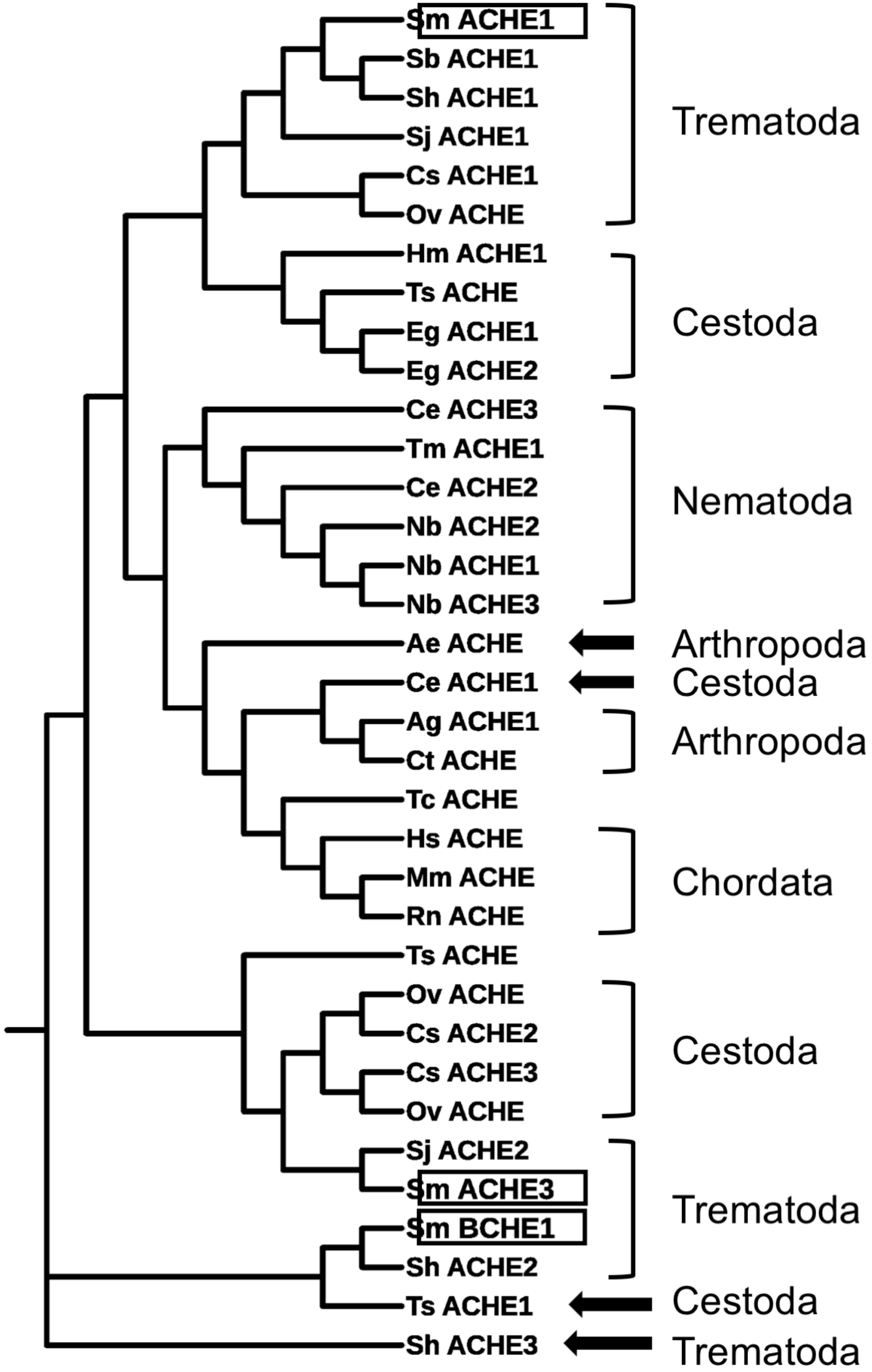

### figure S4

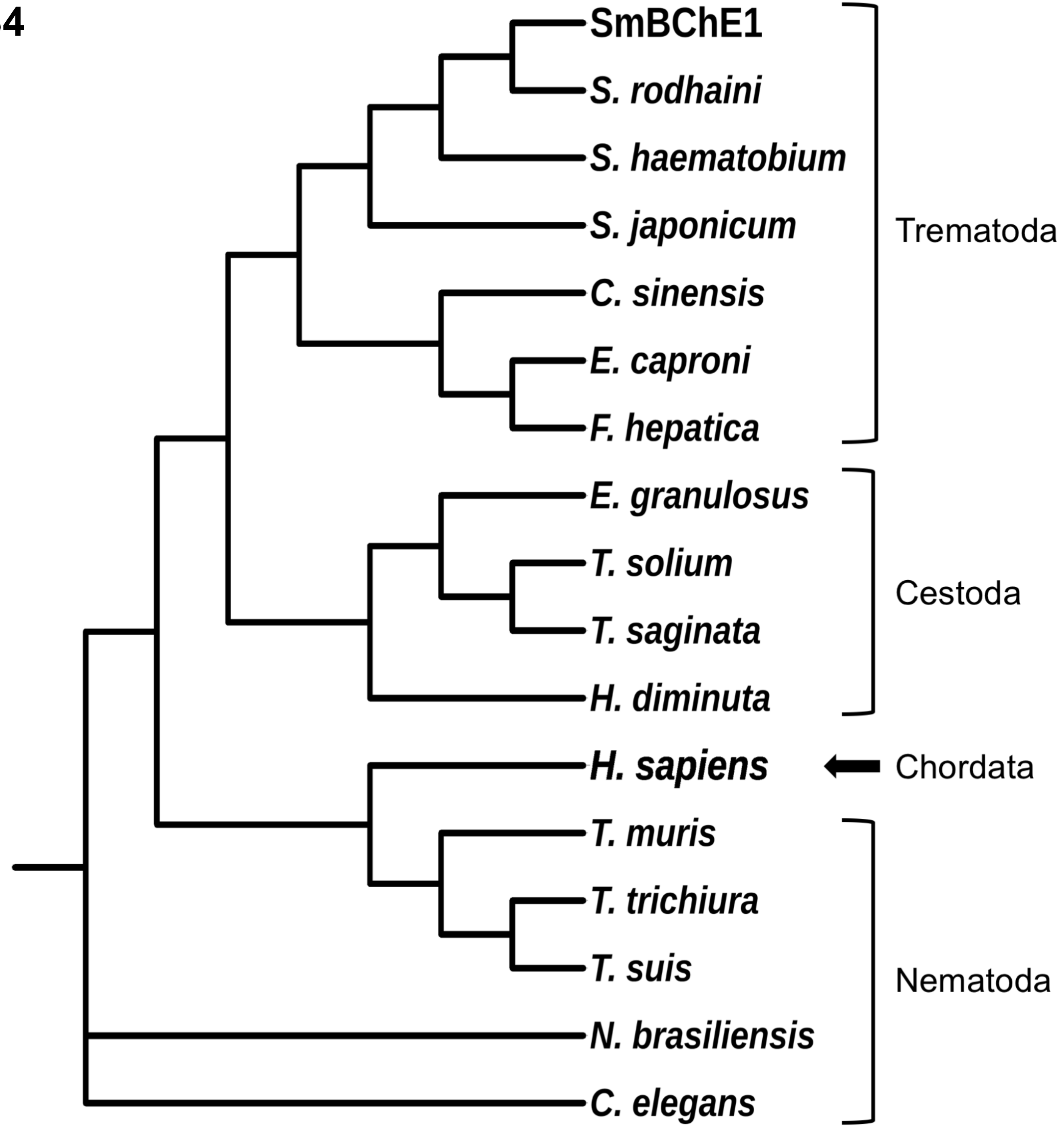

### figure S5

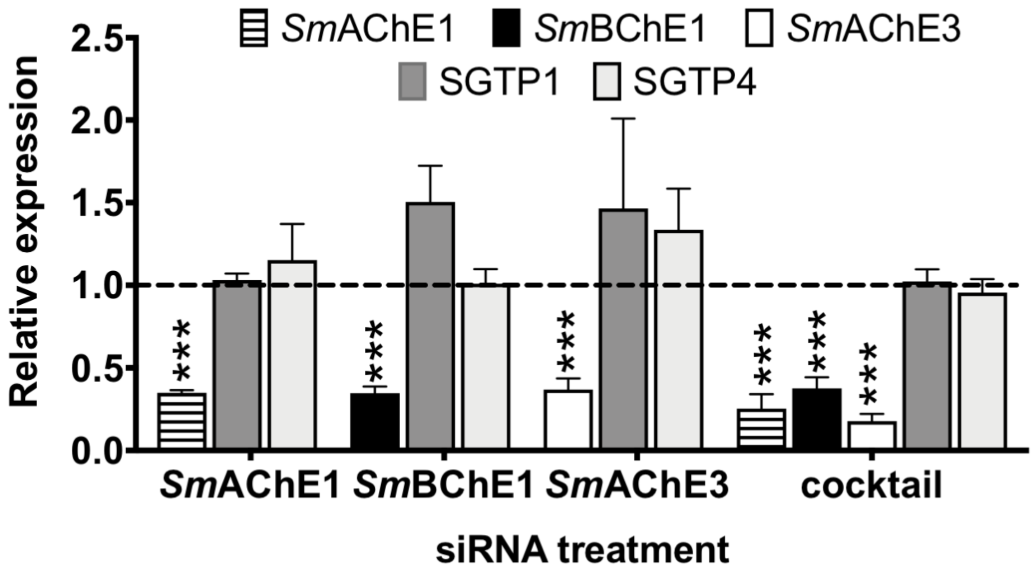
